# Spatial and temporal plasticity of neoantigen-specific T-cell responses bases on characteristics associated to antigen and TCR

**DOI:** 10.1101/2021.02.02.428777

**Authors:** Eva Bräunlein, Gaia Lupoli, Esam T. Abualrous, Niklas de Andrade Krätzig, Dario Gosmann, Franziska Füchsl, Lukas Wietbrock, Sebastian Lange, Thomas Engleitner, Huan Lan, Stefan Audehm, Manuel Effenberger, Melanie Boxberg, Katja Steiger, Yinshui Chang, Kai Yu, Cigdem Atay, Florian Bassermann, Wilko Weichert, Dirk H. Busch, Roland Rad, Christian Freund, Iris Antes, Angela M. Krackhardt

## Abstract

Neoantigens derived from somatic mutations have been demonstrated to correlate with therapeutic responses mediated by treatment with immune checkpoint inhibitors. Neoantigens are therefore highly attractive targets for the development of personalized medicine approaches although their quality and associated immune responses is not yet well understood. In a case study of metastatic malignant melanoma, we performed an in-depth characterization of neoantigens and respective T-cell responses in the context of immunotherapy with Ipilimumab. Three neoantigens identified either by immunopeptidomics or in silico prediction were investigated using binding affinity analyses and structural simulations. We isolated seven T-cell receptors (TCRs) from the patient immune repertoire recognizing these antigens. TCRs were compared in-vitro and in-vivo with multi-parametric analyses. Identified immunogenic peptides showed similar binding affinities to the human leukocyte antigen (HLA) complex and comparable differences to their wildtype counterparts in molecular dynamic simulations. Nevertheless, isolated TCRs differed substantially in functionality and frequency. In fact, TCRs with comparably lower functional avidity and higher potential for cross-reactivity provided at least equal anti-tumor immune responses in vivo. Of note, these TCRs showed a reduced activation pattern upon primary in vitro stimulation. Exploration of the TCR-β repertoire in blood and in different tumor-related tissues over three years, offered insights on the high frequency and particular long-term persistence of low-avidity TCRs. These data indicate that qualitative differences of neoantigen-specific TCRs and their impact on function and longevity need to be considered for neoantigen targeting by adoptive T-cell therapy using TCR-transgenic T cells.

**Statement of translational relevance:** Immunotherapy has demonstrated high efficacy in diverse malignancies. Neoantigens derived from mutations provide promising targets for safe and highly tumor-specific therapeutic approaches. Yet, single determinants of an effective and enduring T-cell mediated tumor rejection are still not well understood. We analyzed in detail seven neoantigen-specific T-cell receptors (TCRs) derived from a melanoma patient targeting three different altered peptide ligands identified by mass spectrometry and prediction analyses. Functional characterization of these TCRs revealed potent anti-tumor reactivity of all TCRs. Of special interest, TCRs with comparably lower affinity demonstrated effective in vivo activity as well as dominant spatial and temporal distribution in blood and tissue. Functional differences of TCR may require further T-cell and/or TCR engineering and should be considered for future clinical trial designs.

## Introduction

Cancer immunotherapy has demonstrated high efficacy in the treatment of diverse malignant diseases as shown by the efficacy of immune checkpoint modulation (1). These therapies can unleash anti-tumor immune responses, thereby conferring recognition and eradication of cancers by the patients’ own immune system without preexisting definition of targets or effector T-cell specificity. However, not all patients benefit from this therapeutic approach. Thus, understanding the nature of tumor recognition versus escape remains of fundamental importance to develop novel immunotherapeutic approaches and improve patients’ outcomes. The tumor mutational load is an important prognostic and predictive biomarker for therapeutic efficacy of immune checkpoint inhibitors across various disease entities, emphasizing the role of neoepitopes in tumor recognition and rejection (2). Therefore, identification and characterization of neoantigens are of particular interest for generation of new personalized immunotherapies, such as vaccination and cellular therapies (3-5). In order to fully exploit the potential of neoantigen-directed immunotherapy, more efforts are needed to understand qualities of neoepitopes and respective T-cell receptors (TCRs). We believe that case studies are currently a valid source to investigate the multitude of aspects involved (6) and that more detailed and comprehensive data are necessary to this purpose. We have previously identified two neoantigens by mass spectrometry (MS) and validated these by proof of defined neoantigen-specific autologous T-cell responses in a patient with melanoma (7). We have now extended the number of immunogenic antigens in this patient using additional prediction analyses and characterized neoantigens by 3D modeling as well as experimental validation of MHC-peptide stability. In addition, we identified seven TCRs recognizing these antigens. These TCRs were monitored in blood and different tissues of the patient over time and functionally characterized in vitro as well as in vivo providing evidence of substantial differences regarding functionality and longevity to be considered for further comprehensive immunotherapies.

## Materials and Methods

### Study approval

Informed consent of all healthy donors and the patient was obtained following requirements of the institutional review board (Ethics Commission, Faculty of Medicine, TU München) and in accordance with principles of the declaration of Helsinki.

### Primary human specimens and cell lines

An overview on clinical course of melanoma patient Mel15 is given in Supplementary Fig. S1, as well as in Supplementary Material and Methods in more detail. All resected samples were handled as previously described for confirmation of diagnosis and expansion of tumor infiltrating lymphocytes (TILs) from a lung metastasis (M_Lung_) (7). Peripheral blood mononuclear cells (PBMCs) were isolated by density-gradient centrifugation (Ficoll-Hypaque, Biochrom) immediately upon receipt using EDTA-anticoagulated blood from Mel15 and blood or apheresis products from healthy donors followed by storage in liquid nitrogen.

PBMCs and T cells used for experiments were cultivated as previously described (7). CD8^+^ T cells used for transduction were obtained by magnetic separation from healthy-donor derived apheresis products (Dynabeads® Untouched™ Human CD8^+^ T Cells Kit, Thermo Fisher; Magnet, DynaMag™).

Cell lines used in this study are: T2 (ATCC Cat# CRL-1992, RRID:CVCL_2211) purchased from ATCC in 2005; lymphoblastoid cell lines HOM-2 (ECACC Cat# 98092902, RRID:CVCL_A612), SWEIG007 (ECACC Cat# 88052037, RRID:CVCL_E838), AMALA (ICLC Cat# HTL14002, RRID:CVCL_E456), OZB (ECACC Cat# 94022545, RRID:CVCL_E799), RSH (ECACC Cat# 88052021, RRID:CVCL_E821), KLO (ECACC Cat# 94050324, RRID:CVCL_E727), LWAGS (ECACC Cat# 88052078, RRID:CVCL_E762), BM21 (ECACC Cat# 88052043, RRID:CVCL_E488), kindly provided by Steve Marsh in 2007; human metastatic melanoma cell line A2058 (ECACC Cat# 91100402, RRID:CVCL_1059, human colon carcinoma cell line MDST8 (ECACC Cat# 99011801, RRID:CVCL_2588) purchased from Sigma-Aldrich in 2018; B-cell lymphoma cell line U-698-M (DSMZ Cat# ACC-4, RRID:CVCL_0017) acquired from DSMZ in 2018. Lymphoblastoid cell line (LCL8) was generated in our laboratory by infection and immortalization of healthy donor-derived B cells with Epstein Barr-virus (EBV)-containing supernatant in 2011. Absence of mycoplasma infection in cell line cultures was routinely confirmed by PCR (Venor GeM mycoplasma detection kit, Minerva Biolabs). Culture conditions are described in Supplementary Material & Methods.

### DNA extraction, exome and TCR-β sequencing

For exome and TCR-beta (TCR-β) deep sequencing, genomic DNA (gDNA) was extracted from a lung biopsy (B_Lung_), an intestinal metastasis (M_Int_), M_Lung_ and draining lymph nodes (LNs) after microdissections of respective regions of interest. The extraction of nucleic acids was performed on 2 μm formalin-fixed, paraffin-embedded (FFPE) tissue slides using Maxwell® RSC Blood DNA Kit and Maxwell® RSC RNA FFPE Kit (Promega) following the manufacturer’s recommendations. DNA was extracted from PBMCs obtained from diverse blood withdrawals (Supplementary Fig. S1) and M_Lung_-derived TILs with DNeasy Blood & Tissue Kit (Qiagen). Whole exome sequencing and single nucleotide variant analyses from M_Int_ were available from our previous publication and were afterwards obtained for M_Lung_, as previously described (7). Next-generation sequencing (NGS) of TCR-β loci was performed by Adaptive Biotechnologies with ImmunoSEQ™ platform at the deep level (exception made for B_Lung_ gDNA which underwent TCR-β sequencing survey level only due to limited material).

### In silico prediction of peptide ligands and HLA binding affinity

Putative mutated nonamer peptide ligands originating from SNVs, identified through a stringent variant calling approach (7), were predicted by translating all sequences bearing a mutation to 23-residue-long amino acid (aa) sequences (mutated aa in 12^th^ position). Protein transcripts were downloaded from Ensembl GRCh38, release 86. In cases where the mutation was located less than 12 aa from the 3’ or 5’ terminus of the gene, the string was shorter and the mutation not centrally located. Peptide-HLA class I binding affinity was predicted for Mel15 HLA alleles HLA-A03:01 and HLA-B27:05 using NetMHC 4.0. Nonamer peptides were ranked by predicted affinity (cutoff < 500 nM) (Supplementary Tables S1 and S2). IC_50_ in-vitro measurements were performed as described in Supplementary Material & Methods.

### Immunogenicity assessment of mutated peptide ligands

For immunogenicity assessment, peptide ligands were ordered from Genscript and DgPeptides. Ligands identified by mass spectrometry, including NCAPG2^P333L^ and SYTL4^S363F^, were tested for recall antigen-experienced T-cell responses as described in Supplementary Material and Methods. Predicted peptides were arranged in pools according to predicted affinity (Supplementary Table S3) and in vitro screened for immunogenicity following protocols described in Supplementary Materials and Methods. Reactivity was assessed by specific IFN-γ release by ELISpot assay by coating ELISpot plates (MAHAS4510) with IFN-γ capture antibody (MABTECH Cat# 3420-3-250, RRID:AB_907283) followed by application of IFN-γ-detection antibody (MABTECH Cat# 3420-6-250, RRID:AB_907273) and visualization by Streptavidin horseradish peroxidase (HRP) (MABTECH Cat# 3310-9-1000).

### Structural modeling of neoantigen binding to respective HLA complexes

Neoantigens and wild-type (WT) peptides were modelled on experimental structures of HLA-A03:01 (PDB-ID: 2XPG (8)) and HLA-B27:05 (PDB-ID 1W0V (9)) crystallized with 9 amino acid long ligands. Peptides from the template were mutated into neoantigen/WT sequences using IRECS program (10).

Surface accessible area (SASA) values were calculated using the VMD software (version 1.9.2) (11,12) applying a probe radius of 1.4 Å. Further information is provided in Supplementary Materials and Methods. Details about heat-up parameters are provided in Supplementary Table S4.

### Isolation of neoantigen-specific T cells

For isolation of KIF2C^P13L^-specific TCRs, peptide-reactive T-cell lines from day 925 and 945 were expanded in culture for two weeks with IL-7 and IL-15 (5ng/mL; Peprotech) and cultured for 24h with irradiated (100Gy) KIF2C^P13L^-pulsed T2 cells in the presence of IL-7. Afterwards, cells were functionally sorted with CD137 MicroBead Kit (MACS^®^) and cloned by limiting dilutions in the presence of irradiated feeder cells (30Gy), 50U/mL IL-2 (Peprotech) and 30ng/mL OKT-3 (kindly provided by Elisabeth Kremmer). The NCAPG2^P333L^-specific T-cell line derived from day 546 (7) was specifically enriched by multimer-based sorting, cloned by limiting dilution and further expanded. SYTL4^S363F^-specific TCRs were isolated either from pre-stimulated peripheral blood (day 740) or from ex vivo expanded TILs (7) using CD137-based functional enrichment or multimer-based sorting, respectively. Proliferating T-cell clones were screened for IFN-γ secretion using the IFN-γ ELISA Kit II (BD Biosciences) upon co-culture with peptide-pulsed T2 cells. mRNA from reactive clones was extracted with TRIzol™ Reagent (Invitrogen) and transcribed to complementary DNA with Affinity Script Multiple Temperature cDNA Synthesis Kit (Agilent Technologies).

### Molecular cloning and retroviral transduction of TCRs, minigenes and HLA

TCR repertoire PCR analysis followed by Sanger sequencing was performed to determine variable alpha and beta chain usage of neoantigen reactive T-cell clones (13,14). V-D-J rearrangements of complementarity-determining regions (CDR3) were investigated using IMGT/V-Quest (http://www.imgt.org/IMGT_vquest/vquest) and the complete sequences were in silico reconstructed with Ensembl database. TCR sequences were modified by in silico murinization, insertion of an additional cysteine bridge and codon optimization (BioCat) (15,16). Optimized TCR constructs were retrovirally transduced in CD8^+^ healthy donor derived T cells as described before and expanded for 7 to 10 days with IL-7 and IL-15 (5ng/mL; Peprotech) before functional characterization (13,14).

SYTL4 and NCAPG2 mutated and WT minigenes were cloned as previously described (7). KIF2C minigenes were in silico designed, comprising 100 bp up- and downstream of mutated position, synthetized (Genscript) and cloned into pMP71 backbone. Additionally, tandem constructs were in silico designed, synthetized (Genscript) and cloned into the pMP71 backbone with single or tandem constructs for minigenes containing mutated sequences of all three neoantigens and WT counterparts separated in two different vectors. All minigene vectors included a reporter gene dsRed ExpressII to allow sorting of transgenic cells. Neoantigen-coding and WT counterpart minigenes were transduced in different cell lines as LCL-1, U-698-M, A2058 and MDST8 as previously described (7) with small changes for adherent cell lines omitting RetroNectin (Clontech). LCL-1 was transduced with single minigenes, while other cell lines with tandem constructs, followed by single clone selection.

TCR beta and alpha chains as well as tandem constructs of minigenes were separated by a self-cleaving P2A element (13). All vectors were amplified using NEB® 5-alpha Competent E. coli (New England BioLabs) and purified with NucleoBond® Xtra Midi/Maxi (Macherey-Nagel). T2 were retrovirally transduced with the HLA restriction elements HLA-A03:01 and B27:05 as described before (13). Details on cell cultivation are described in Supplementary Material and Methods.

### Flow-cytometry-based assessment of TCR expression

Transduction efficiency was determined by staining of transgenic TCR using an antibody specifically binding to murine TCR constant beta chain (TCRmu-antibody; (clone H57-597) Thermo Fisher Scientific Cat# 11-5961-81, RRID:AB_465322) based on the isotype control (Thermo Fisher Scientific Cat# 11-4888-81, RRID:AB_470037). For multimer staining, Strep-tagged mutated and WT pHLAs (e.g. SYTL4^S363F^ and SYTL4^WT^) were multimerized on a fluorophore labeled Strep-Tactin backbone (Iba GmbH) in a 1 µg : 1 µl ratio. Transduced T cells were stained with multimers, anti-CD8 (BD Biosciences Cat# 557085, RRID:AB_396580 and BD Biosciences Cat# 555634, RRID:AB_395996), anti-CD3 (BD Biosciences Cat# 557694, RRID:AB_396803) and 7-Aminoactinomycin D (7-AAD). All flow cytometry measurements were performed with LSRII flow cytometer (BD Biosciences) and analyzed with FlowJo Software (version 10).

### Functional analyses of TILs and TCR-transgenic effector T cells

TCR-transduced T cells were co-incubated with target cells (LCL-1) transduced with different minigene constructs or pulsed with defined peptides (1 μM). WT minigenes and/or irrelevant peptides served as controls for TCR specificity. Assays for detection of cytokine secretion were performed in triplicates (E:T = 1:1; 10,000 target and effector cells per well). Supernatants from T-cell cultures were used for assessment of a diverse cytokine panel with MACSPlex 12 Cytokines Kit (Miltenyi Biotech) following the manufacturer’s recommendations. Reactivity of TILs was assessed by IFN-γ ELISpot assay on a coculture with autologous peptide-pulsed γ-irradiated LCLs (1µM) for 24 hours using at E:T of 1:1.

Functional avidity was assessed by incubating transgenic T cells with T2 target cells (ATCC® CRL-1992™) pulsed with titrated peptide concentrations (E:T = 1:1; 10,000 cells/well). IFN-γ secretion was quantified by ELISA and values obtained were fitted into a nonlinear variable-slope regression curve on GraphPad Prism 7.

Functional avidity assays were performed at least three times using at least two different transductions and donors showing comparable results.

In-vitro target cell killing was assessed by impedance-based xCELLigence assays as described in Supplementary Material and Methods.

TCR cross-reactivity was tested by stimulation of TCR-transgenic T-cell populations with T2 cells pulsed with ala/thr substitution variants of the three neoepitopes, in direct comparison to the original mutated identified peptides. IFN-γ values from every condition were normalized on the positive control using the following formula:

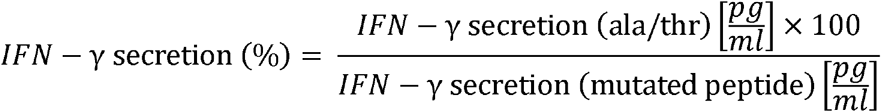

Relative IFN-γ-secretion after incubation with an ala/thr variant above 50% were considered as a replaceable position.

For evaluation of general alloreactivity, TCR-transduced T cells were stimulated with respective neoantigens presented in the context of different HLA class I alleles as well as non-pulsed target cells. Effector cells were cultured with LCL presenting a broad variety of HLA alleles (Supplementary Table S5), pulsed with mutated peptides (E:T = 1:1; 10,000 cells/well). As readout, IFN-γ production was quantified by ELISA.

### In vivo TCR rejection potential

NOD.CG-Prkdc^scid^ IL2rg^tm1Wjl^/SzJ (NSG; The Jackson Laboratory) were maintained according to the institutional guidelines and approval of local authorities. A xenograft murine model was generated as previously described (13,17). NSG mice were subcutaneously injected with U-698-M cells (10×10^6^ cells/flank) transduced with tandem minigene construct coding for defined neoantigens. Growth was monitored in vivo by external measurements with digital caliper. As tumors reached an area of ca. 20 mm^2^, T cells transduced with TCR KIF2C-PBC1, KIF2C-PBC2, SYTL4-PBC1 and SYTL4-TIL1 were injected intravenously. 2×10^7^ T cells were administrated to 5 mice per group (n = 5). Tumor growth kinetics were monitored for 10 days with digital caliper. On day 11, remaining tumors and spleen were excised and passed through a cell strainer. Blood samples were taken and anticoagulated with EDTA. Ammonium-Chloride-Potassium (ACK) lysis (Life Technologies) was additionally performed on spleen and blood samples. Single cell suspensions were stained for detection of transgenic T cells by flow cytometry as previously explained.

### Statistical analysis

Significance of differences within half maximal effective concentration (EC_50_) values of analyzed TCRs were investigated by one-way ANOVA and Tukey’s multiple comparison test. With regard to in-vivo rejection potential of the TCRs, differences in tumor growth were calculated with two-way ANOVA test (time; treatment) and Dunnett’s test for multiple comparisons. Flow cytometry results from in-vivo experiment were evaluated with Kruskal-Wallis test and uncorrected Dunn’s test for multiple comparisons. To calculate the statistical significance of the increase in SASA, a standard independent two-sample t-test was used. Statistical analyses were performed with GraphPad Prism 7.04 software.

Additional Information on Material and Methods is provided in the Supplemental file.

## Results

We have previously reported mass spectrometry (MS)-based identification of two immunogenic neoantigens (SYTL4^S363F^ and NCAPG2^P333L^) presented by an intestinal metastasis (M_Int_) of a melanoma patient (7). Based on this case we performed multi-dimensional analyses using a comprehensive material collection including multiple additional biopsies and resected tumor material (B_Lung_, M_Lung_, M_Int_-LN1, M_Int_-LN2 and M_Lung_-LN), as well as blood specimens, collected over a period of 3 years. The disease course of this patient is shown in Supplementary Fig. S1.

### In-silico predictions complement MS-based neoantigen identification

In order to investigate whether critical neoantigens may have been missed by MS, a sequence-based prediction approach was applied. 1,196 missense mutations were called from M_Int_-tumor tissue, leading to prediction of ∼4670 peptides (8-12 aa long) with binding affinity < 500nM. By sorting nine-aa-long putative peptides according to NetMHC predicted binding affinity, previously identified neoantigens NCAPG2^P333L^ and SYTL4^S363F^ ranked 24^th^ and 6t^th^ in HLA-A03:01 and HLA-B27:05 lists respectively (Supplementary Tables S1 and S2). For most peptides, high binding affinities could be confirmed experimentally, however, binding affinities partially differed substantially to predicted ones (Supplementary Tables S1 and S2). Investigation of immunogenicity of these top ranked nonamers binding to HLA-A03:01 and B27:05 alleles resulted in the detection of one additional confirmed neoepitope, KIF2C^P13L^ (Supplementary Fig. S2; Table 1), ranking position 18^th^ of top 25 HLA-A03:01 binding peptides according to NetMHC 4.0 (Supplementary Table S1). Molecular dynamic (MD) simulations of MHC peptide binding of SYTL4^S363F^, KIF2C^P13L^ and NCAPG2^P333L^ were subsequently performed in order to investigate differences in solvent accessible surface areas. Clear differences between wildtype and mutated peptides were observed (Supplementary Fig. S3). This was even the case for the mutated P2 of NCAPG2^P333L^ harboring the mutation inside the peptide binding cleft and not at the TCR interface (Supplementary Fig. S3C) as the larger size of the mutated residue (Leu => Pro) shifted the position of solvent-exposed P4 Leu, increasing its SASA significantly from 108.06 Å^2^ (WT) to 133.54 Å^2^ (mutant).

**Table 1.**
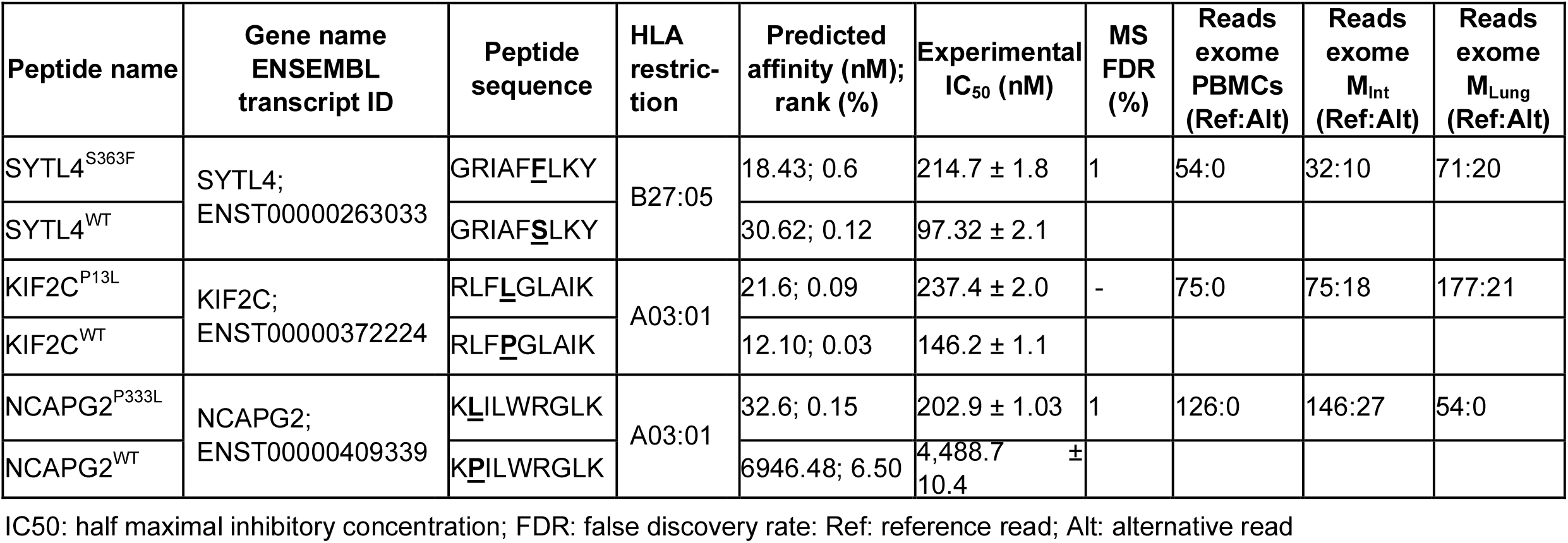
Characteristics of neoantigens targeted by identified TCRs

### In-depth characterization of immune responses against three selected neoantigens show high antigen specificity

Immune responses against KIF2C^P13L^, SYTL4^S363F^ and NCAPG2^P333L^ neoantigens (Table 1) were further analyzed in detail and seven TCRs with specificity to these neoantigens were isolated (Supplementary Table S6). Transduction efficiency by staining of murinized constant regions of transduced TCRs showed differences for defined TCRs, with lowest transduction rates for TCRs KIF2C-PBC1 and SYTL4-TIL2 (Supplementary Fig. S4). Moreover, by performing HLA class I multimer staining for all TCRs, T-cell populations binding the mutated peptide multimer could be detected only for five out of seven TCRs (Supplementary Fig. S5). In-vitro functional characterization of TCR-transgenic CD8^+^ T cells showed highly specific IFN-γ secretion upon recognition of target cells pulsed with mutant peptides or transduced with minigenes coding for neoantigens (Fig. 1A-C). No reactivity was observed against WT or irrelevant peptides as well as the corresponding control minigenes. Of note, transduction rates, evaluated by TCRmu staining (Supplementary Fig. S4), did not necessarily correlate with IFN-γ secretion as both KIF2C^P13L^-specific TCRs showed similar reactivity to LCL-1 presenting the mutant peptide despite major differences in TCR expression (Fig. 1B). In line with that, cytokine secretion of SYTL4^S363F^-specific TCRs was substantially higher in comparison to those reactive to KIF2C^P13L^ or NCAPG2^P333L^, even for those TCRs with lower surface expression (Figs. 1A-C).

**Figure 1.**
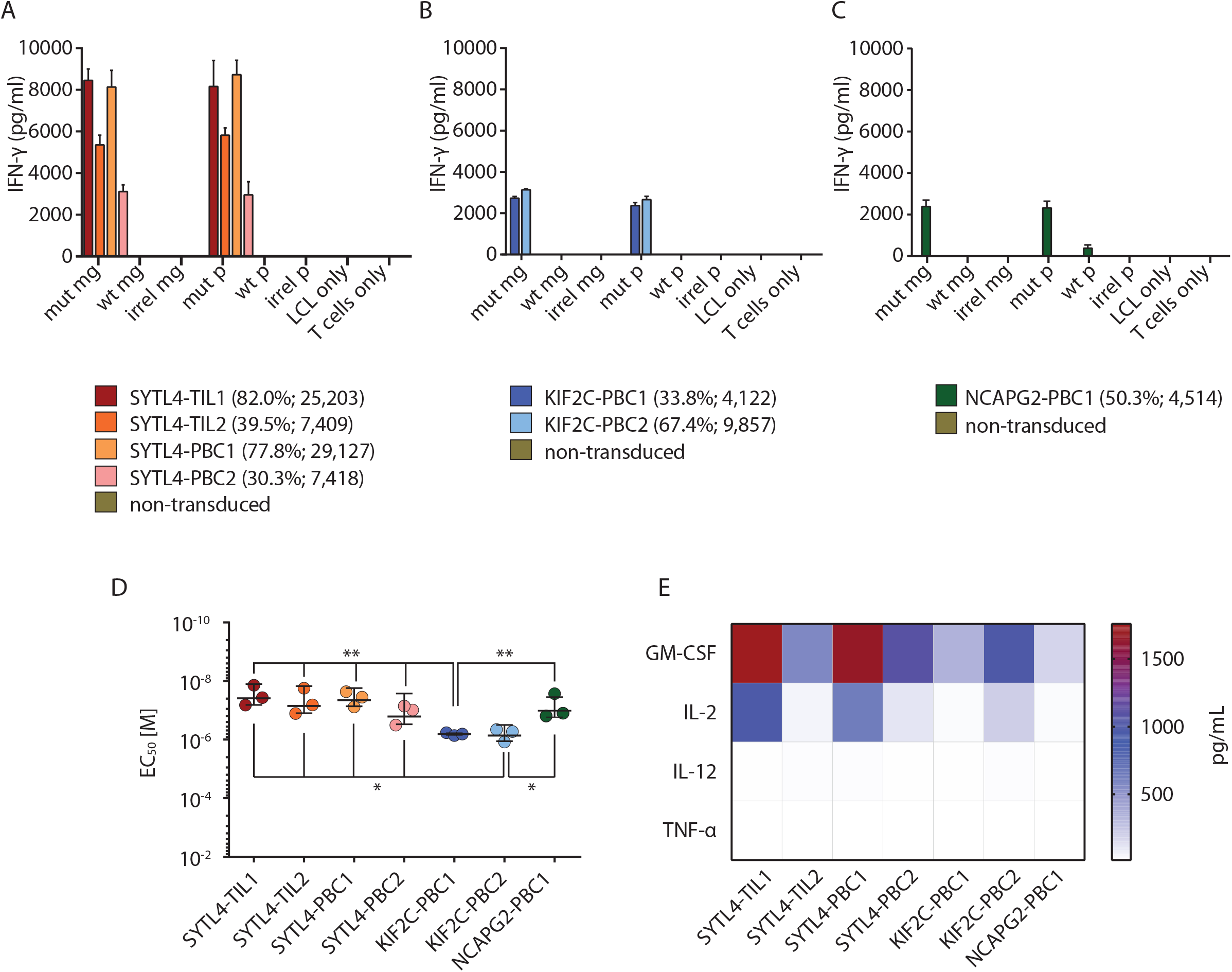
High specificity and strong functional performance of identified TCRs in vitro. **A, B** and **C** secretion of IFN-γ by CD8^+^ T cells transduced with TCRs specific for SYTL4^S363F^ (**A**), KIF2C^P13L^ (**B**) and NCAPG2^P333L^ (**C**) after coculture with LCL-1 presenting neoantigens and respective WT counterparts at E:T = 1:1 is shown. LCL-1 transduced with minigenes (mg) encoding for fragments of mutant sequence (mut mg), WT sequence (WT mg) or irrelevant sequence (irrel mg) were compared to peptide pulsed LCLs (mut p, WT p, irrel p; 1 μM peptide). IFN-γ secretion in supernatants was investigated by ELISA assay. Bars represent average reads from three duplicates, error bars represent SD. Transduction efficiencies and MFI values are indicated below the graphs. **D**, Comparison of functional avidity of neoantigen-specific TCRs, calculated as EC_50_ of cognate mutated peptide. IFN-γ secretion was assessed on supernatants and a non-linear curve was fit to determine the EC_50_ value. EC_50_ values deriving from three different experiments were depicted for each TCR. Bars in the graph represent the mean value and SD. Significance is calculated with one-way ANOVA and Tukey multiple comparison test (* p ≤ 0.05, ** p ≤ 0.01). **E**, Assessment of multi-cytokine secretion of TCR-transduced T cells upon coculture with LCL-1 pulsed with mutated peptides. All experiments were performed at least with three different sets of transduced T cells derived from two different healthy donors.

Functional avidity measurements using peptide-pulsed T2-A3 and T2-B27 cells showed EC_50_ values against respective ligands in the almost nanomolar range for all TCR-transduced T cells (Fig. 1D). However, TCRs specific for SYTL4^S363F^ and NCAPG2^P333L^ showed significantly higher avidities compared to KIF2C^P13L^-specific TCRs (Fig. 1D, Supplementary Fig. S6).

To further decipher functional differences between all seven TCRs, defined cytokines as GM-CSF, IL-2, IL-12 and TNF-α, were analyzed by multiplex analysis using supernatants of TCR-transgenic CD8^+^ T cells stimulated with mutated or WT minigene-transduced LCL-1 target cells (Fig. 1E). T cells transduced with TCRs SYTL4-TIL1 and SYTL4-PBC1 secreted highest concentrations of these cytokines in response to respective mutated minigenes.

To analyze the dynamics of TCR-mediated cytotoxic activity in vitro, an impedance-based xCELLigence system was used to monitor growth kinetics of minigene transduced target cell lines after co-incubation with TCR-transduced T cells. With the exception of SYTL4-PBC2, all TCRs mediated 100% of lysis of MDST8^MUT^ and A2058^MUT^ target cells within 12-16h while WT clones were not affected (Supplementary Fig. S7A-C). Among SYTL4^S363F^-specific TCRs, SYTL4-TIL1 showed the fastest rejection dynamics (Supplementary Fig. S7A). With respect to KIF2C^P13L^-specific TCRs, rejection dynamics again appear to be similar despite different transduction rates (Supplementary Fig. S7B). Comparison of cytotoxic potential for all TCRs showed different dynamics dependent on the target cells but comparable final effects except for SYTL4-PBC2 displaying an inferior cytolysis pattern (Supplementary Fig. S7D).

### TCRs show neoantigen-specific binding and low cross-reactivity patterns

A set of altered peptide ligands containing individual alanine or threonine replacements (ala/thr scan) at every single position of the cognate neoantigen was used to investigate the cross-reactivity potential of defined TCR-transgenic T cells. Highly similar recognition patterns of all four SYTL4^S363F^-specific TCRs were observed despite highly diverse TCR sequences (Fig. 2A; Supplementary Table S7). These results suggest a similar peptide-HLA docking pattern for all TCRs with this neoantigen specificity. In contrast, recognition patterns of exchange ala/thr motifs were slightly more variable with respect to both KIF2C^P13L^-specific TCRs (Fig. 2B). This was reflected by a higher potential for cross-reactivity against naturally occurring peptides within the human proteome, as investigated by ScanProsite tool (https://prosite.expasy.org/scanprosite) (18). SYTL4^S363F^-and NCAPG2^P333L^-specific TCRs exhibited potential cross-reactivity with four to six antigens only, while KIF2C^P13L^-specific TCRs might recognize > 400 targets (Supplementary Table S7). Thus, these data suggest a higher probability of cross-recognition of antigens by TCRs specifically recognizing KIF2C^P13L^ in comparison to those specific for SYTL4^S363F^ and NCAPG2^P333L^ (Fig. 2A-C). Otherwise, the alloreactive potential of selected TCRs tested against a panel of LCLs expressing the most frequent HLA allotypes (Supplementary Table S5) with or without previous peptide pulsing showed low alloreactive potential for all neoantigen-specific TCRs (Supplementary Fig. S8).

**Figure 2.**
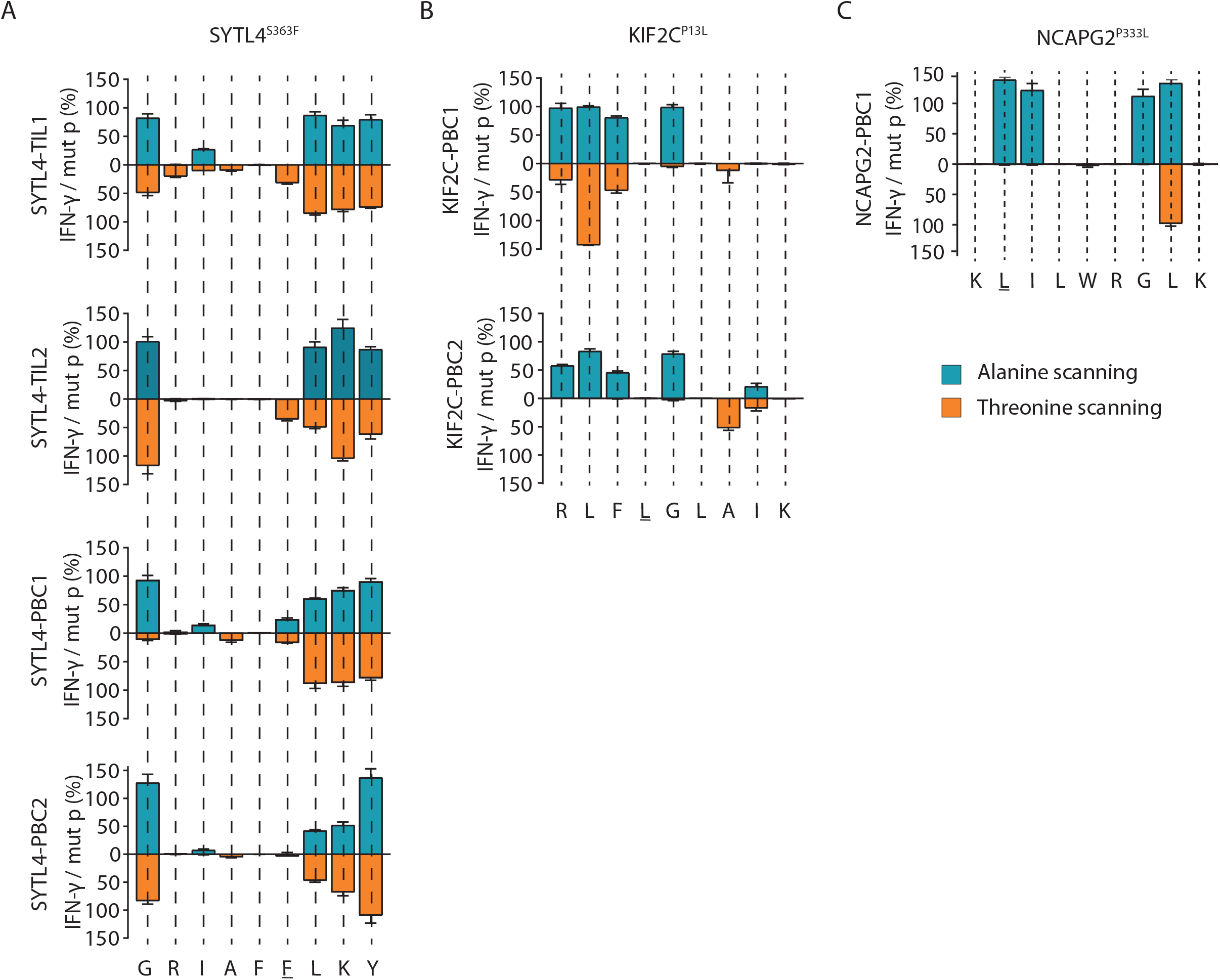
Cross-reactivity analyses indicate antigen-dependent TCR docking on HLA peptide complexes. **A, B** and **C**, TCR cross-reactivity was tested by quantification of secreted IFN-γ upon coculturing TCR-transduced T cells with T2 target cells pulsed with ala/thr scanned peptide cognates (1 μM peptide) of ligands SYTL4^S363F^ (**A**), KIF2C^P13L^ (**B**) and NCAPG2^P333L^ (**C**). IFN-γ secretion values from single conditions were normalized against cytokine level in response to the defined neoantigen.

### Neoantigen-specific TCRs display in vivo anti-tumor potential

We next asked, whether differences of in-vitro functionality especially with respect to SYTL4^S363F^ and KIF2C^P13L^–specific TCR may also lead to different in vivo tumor rejection patterns (Fig. 3). Therefore, anti-tumor reactivity of defined neoantigen-specific TCRs was investigated in a xenograft murine model using a minigene-transduced U-698-M lymphoma model. U-698-M tumor growth in NSG mice was monitored starting from two days before injection of TCR-transduced T cells and non-transduced T cells. Around day 5 post injection, tumors of mice that received non-transduced T cells kept constantly growing, while tumors in hosts receiving T cells transduced with TCRs SYTL4-TIL1, SYTL4-PBC1, KIF2C-PBC1 and KIF2C-PBC2 showed a significant and comparable delay in growth despite low transduction rates (p < 0.001) (Fig. 3A). Flow cytometry data show significant infiltration of mutated tumors by respective TCRs (SYTL4-TIL1: p = 0.02, SYTL4-PBC1: p = 0.02 and KIF2C-PBC2: p = 0.01, (Fig. 3B) as well as presence of TCR-transgenic T cells in the spleen (Fig. 3C).

**Figure 3.**
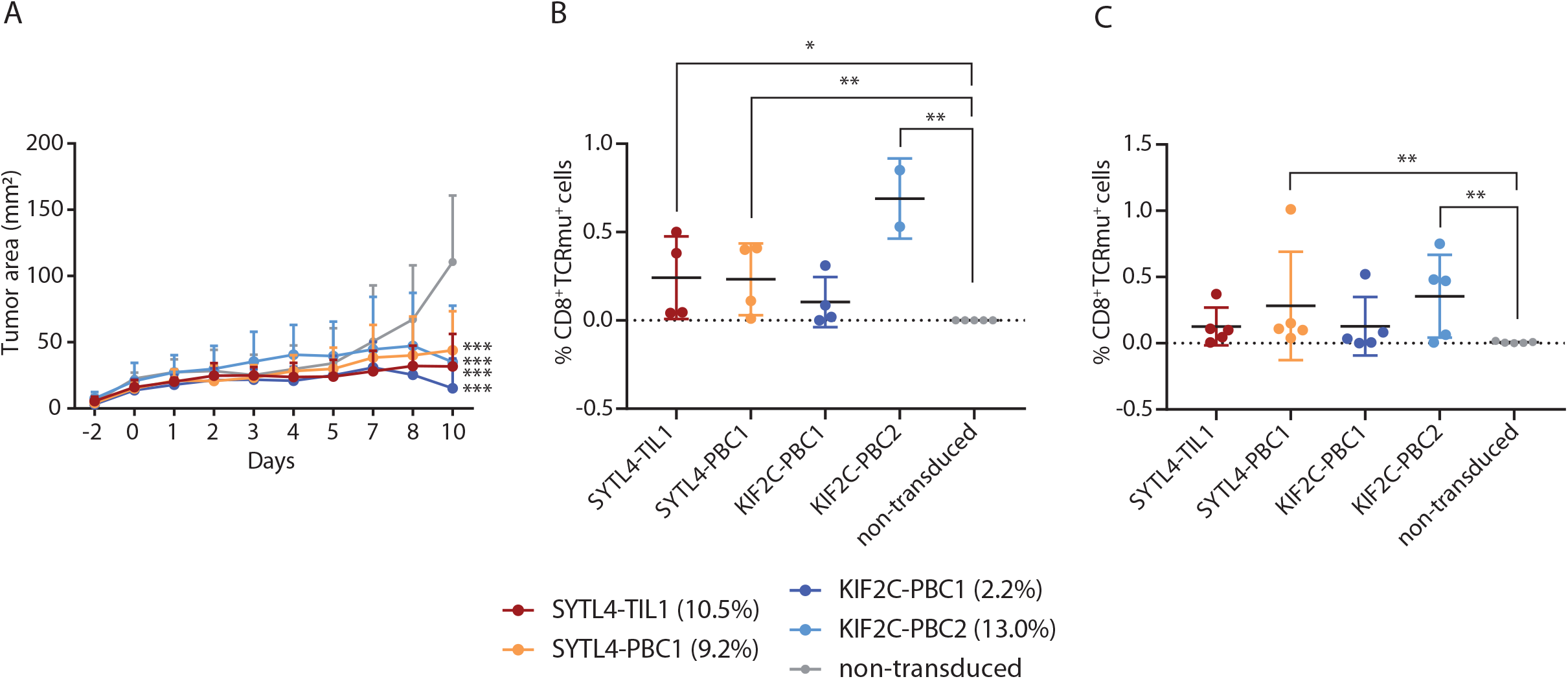
Performance of T cells transgenic for neoantigen-specific TCRs in vivo. **A**, growth kinetics of U-698-M tumors expressing neoantigens (area in mm^2^) in NSG mice. Mean values and SD are depicted for each group of mice bearing tumors (n = 5). Animals were i.v injected with 2×10^7^ T cells on day 0. Significance is calculated with two-way ANOVA (time; treatment) and multiple comparison Dunnett’s test. **B** and **C**; the percentage of CD8^+^ TCRmu^+^ T cells calculated on total alive cells, detected in the tumor (**B**), and spleen (**C**) is depicted. Significance is calculated with Kruskal-Wallis test. TCR transduction efficiencies are indicated below the graphs. * p = 0.033; ** p = 0.002; *** p < 0.001.

### Antigen-dependent spatial and temporal distribution of identified TCRs in blood and tissues

To investigate spatial and temporal distribution of tumor antigen-specific TCR clonotypes within the patient’s tissues and blood, TCR-β deep sequencing was performed by extracting genomic DNA from B_Lung_, M_Int_, M_Lung_, LNs, as well as PBMCs from different time points (Supplementary Fig. S1).

Overlap of TCR-β clonotypes was investigated in all three tumor tissues. All tumor samples, resected over a timespan of eight years, share 29 TCR-derived CDR3 (Supplementary Fig. S9A). M_Int_ and M_Lung_, closer in time, share 3,072 TCR clonotypes (14.77% of all sequences identified in M_Int_ and M_Lung_). Notably, a number of 462 TCR clonotypes was shared between both resected metastases and all analyzed adjacent lymph nodes (Supplementary Fig. S9B). The seven characterized neoantigen-specific TCRs were tracked within metastasized tumor lesions, lymph nodes and blood samples (Fig. 4; Supplementary Fig. S9). Frequencies of these TCRs identified from peripheral blood specimens, lymph nodes and tumor tissue are shown in Supplementary Tables S8 and S9. KIF2C^P13L^-specific TCRs were the most abundant in all metastasized tumor tissues over time and TCR KIF2C-PBC1 could be detected in B_Lung_ even prior to therapy with Ipilimumab. Overall, frequency of KIF2C^P13L^-specific TCRs, especially KIF2C-PBC2, was increasing in M_Lung_ compared to M_Int_ (Fig. 4A; Supplementary Table S8), whereas frequencies of SYTL4^S363F^-specific TCRs were decreasing or remained stable. Notably, TCR NCAPG2-PBC1 could not be detected in M_Lung_ corresponding to the lack of detectability of this specific mutation in the metastasis on exome level (Fig. 4A; Table 1). By investigating the relative frequency of known neoantigen-specific TCRs amongst all identified CDR3 sequences in tumor-associated non-malignant lymph nodes, the overall percentage of all seven cognate TCRs contributing to the analyzed TCR repertoire was calculated ranging from 0.11% to 1.53% (Fig. 4B). Thereby, TCRs specific for KIF2C^P13L^ comprised the highest relative percentage not only in tumor tissues but also in tumor-associated lymph nodes. However, we observed relative frequency of defined TCR clonotypes being more evenly distributed in associated lymph nodes (Fig. 4B).

**Figure 4.**
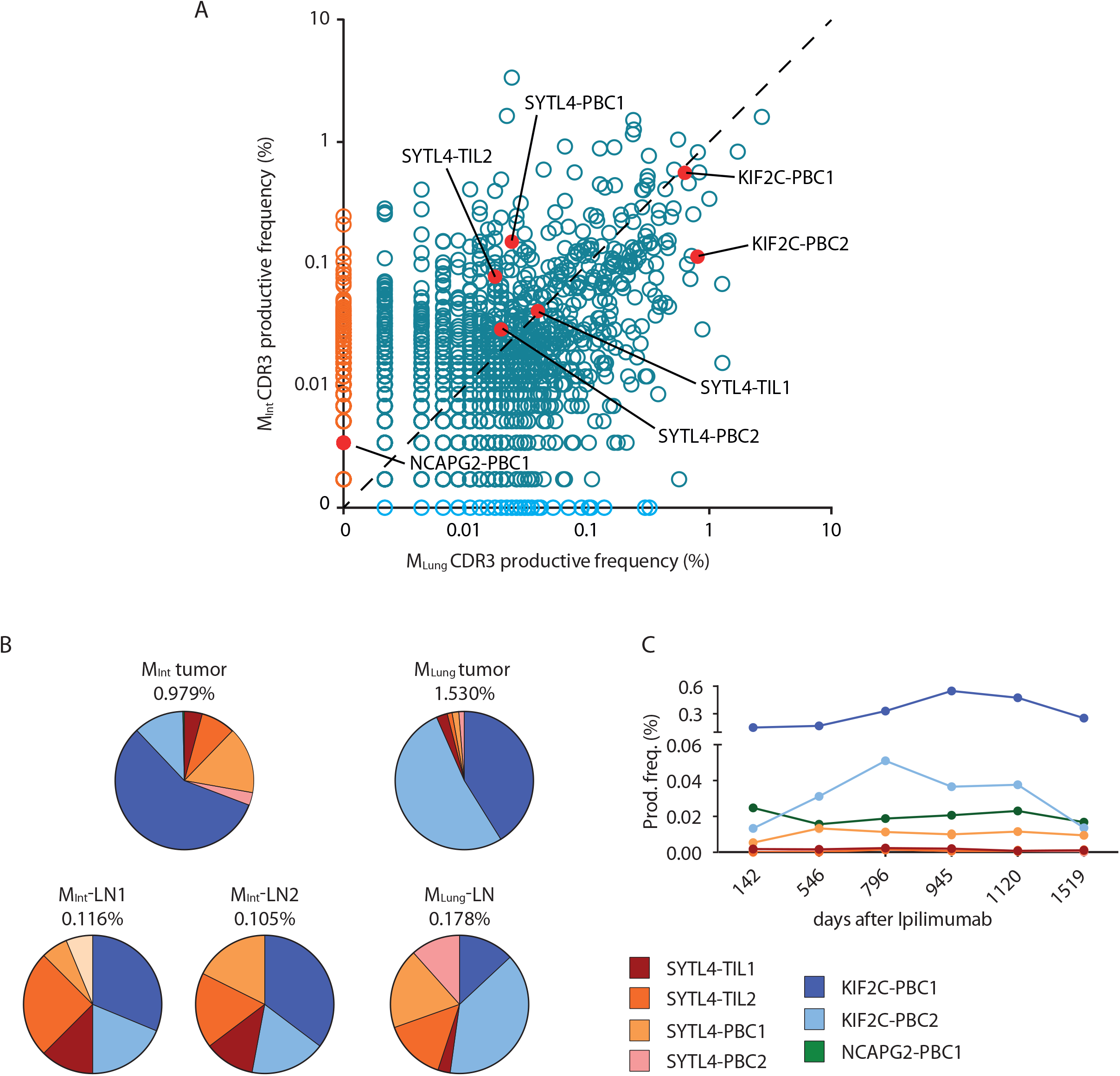
Spatial and temporal distribution of neoantigen-specific TCRs as determined by TCRβ sequencing. **A**, Productive frequency of CDR3 rearrangements (amino acid) calculated as number of sequencing templates divided by the sum of template counts for all productive rearrangements in M_Int_ and M_Lung_. Scatter plot dataset was generated with Adaptive Biotechnologies ImmunoSEQ analysis software. **B**, Distribution of neoantigen-specific TCRs in resected tumor and lymph node tissues. Percentages indicate frequency of identified neoantigen-specific TCRs in analyzed tissue repertoire as inferred from TCR-β deep sequencing data. **C**, Productive frequency expressed as percentage of neoantigen-specific clonotypes in peripheral blood at different time points after first treatment with Ipilimumab.

In peripheral blood, all specific TCRs could be detected in at least four out of the six sequenced samples (Fig. 4C). Productive frequencies of SYTL4^S363F^-specific TCRs ranged between 0.0008% to maximal 0.013%, whereas TCRs KIF2C-PBC1, KIF2C-PBC2 and NCAPG2-PBC1 show considerably higher frequencies between 0.013% and 0.54% in analyzed blood samples (Supplementary Table S9). Of note, TCR KIF2C-PBC1 showed again the highest frequencies over time. In contrast, frequency of TCR KIF2C-PBC2 in peripheral blood increases up to the time point of resection the lung metastasis and decreases thereafter corresponding to the high frequency of this TCR in M_Lung_ and associated lymph node (Fig. 4C). Presence of identified TCR KIF2C-PBC2 within respective primary tumor tissue was verified additionally by TCR imaging using RNA hybridization BaseScope assay on tumor M_Lung_ (Supplementary Fig. S10). In addition, mutated ligand KIF2C^P13L^ was also recognized by isolated TILs of the patient (Supplementary Fig. S11).

### T cells specific for SYTL4^S363F^ and KIF2C^P13L^ display divergent activation and dysfunctional profiles

To further delineate TCR-dependent differences, we aimed to investigate activation and dysfunction parameters of neoantigen-specific stimulated T cells. We exemplarily investigated the T-box transcription factor TBX21 (T-bet) and programmed cell death protein 1 (PD-1) expression on SYLT4-TIL1, -PBC1, KIF2C-PBC1 and –PBC2 transduced T cells after co-culture with minigene-transduced U-698-M. SYTL4^S363F^-specific TCR displayed a higher upregulation of T-bet after 24 hours compared to KIF2C^P13L^-specific constructs with all TCR showing a downregulation of expression to baseline after 96 hours (Supplementary Fig. S12A-B). To minimize potential influence of different transduction rates, T cells transduced with TCRs SYTL4-TIL1 and KIF2C-PBC2 were enriched using iRFP-containing constructs. Using enriched populations, marked differences of IFN-γ secretion between TCRs with defined specificities were observed upon antigen-specific stimulation (Fig. 5A), as shown for non-enriched transduced T cells (Fig. 1A-B). Investigating early T-cell activation, we observed a higher upregulation of T-bet for SYTL4-TIL1-iRFP after 24 hours compared to KIF2C-PBC2-iRFP (Fig. 5B-C, Supplementary Fig. S12C-D). Also, upregulation of PD-1 was higher after 24 hours in SYTL4-redirected T cells in comparison to KIF2C-PBC2 with respect to frequency and MFI of PD-1 positive T cells (Fig. 5D-E). Yet, in-vitro re-challenging of neoantigen-specific T cells after 11 days contrarily revealed an almost equal IFN-γ secretion of both TCRs upon stimulation with U-698-M expressing the mutated minigene (Fig. 5F). In this regard, T-bet and PD-1 expression of KIF2C-PBC2-iRFP was at the same level as SYTL4-TIL1-iRFP (Fig. 5G-J).

**Figure 5.**
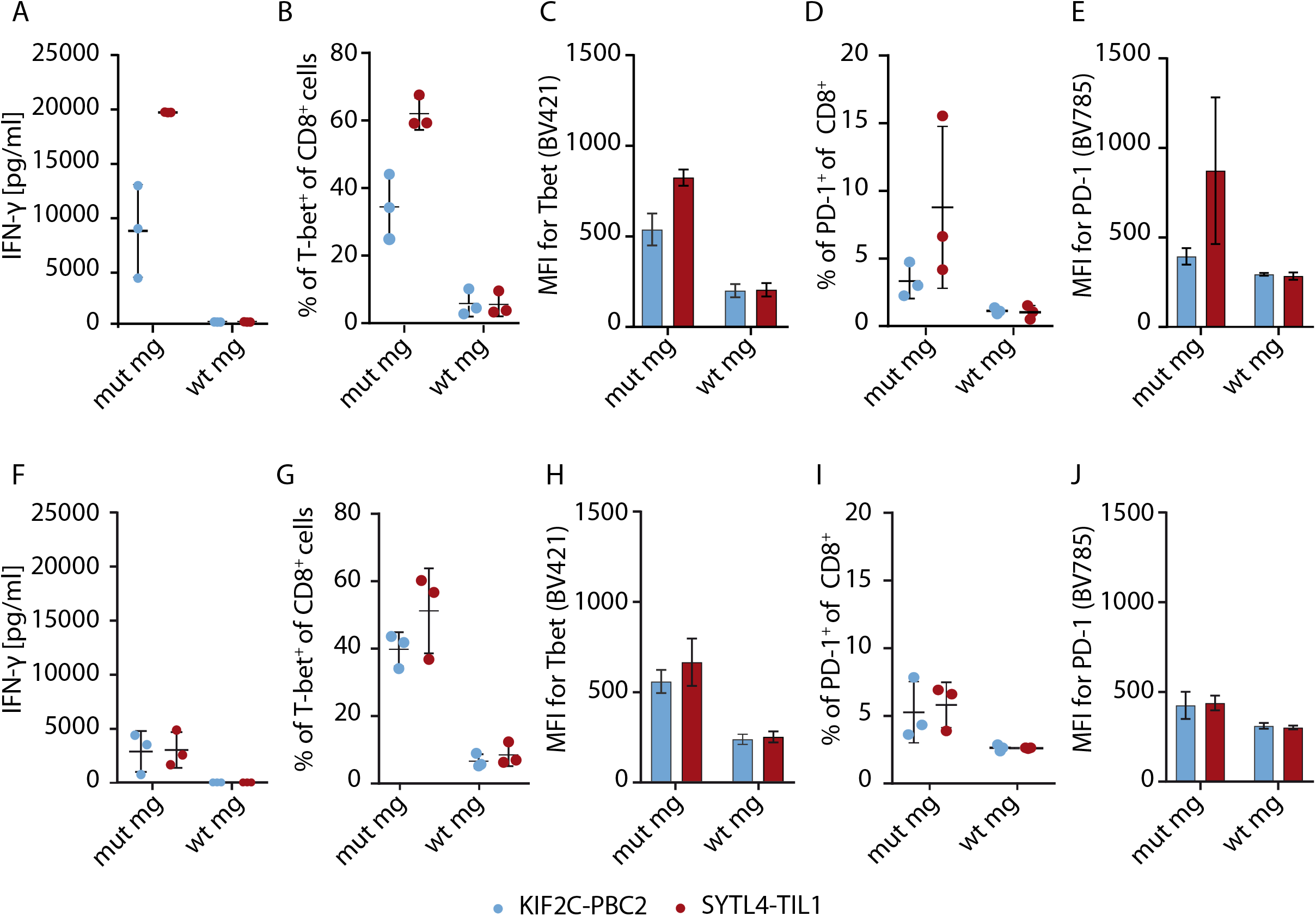
T-cell activation and dysfunction of enriched SYTL4-TIL1-iRFP and KIF2C-PBC1-iRFP TCR upon neoantigen stimulation with minigene-transduced U-698-M. **A**, IFN-γ secretion of T cells 24 hours after coculture. **B-E**, Expression of specific markers 24 hours after coculture: percentage of T-bet positive cells within CD8^+^ T cells (**B**), MFI for T-bet of CD8^+^ T cells (**C**), percentage of PD-1 positive cells within CD8^+^ T cells (**D**) and MFI for PD-1 of CD8^+^ T cells (**E**). **F-J**, T-cell activation and dysfunction was comparably assessed 24 hours after second stimulation of SYLT4-TIL1-iRFP and KIF2C-PBC2-iRFP transgenic T cells with minigene-expressing target cells at day 11 after initial stimulation. Results of transduced T cells from three different donors are shown.

## Discussion

Within this study, we performed an in-depth characterization of TCR specifically recognizing neoantigens derived from somatic mutations which have been identified in malignant tissue of a melanoma patient. We thereby reveal important insights highly relevant for the development of advanced personalized immunotherapies targeting neoantigens.

For neoantigen identification, MS-based immunopeptidomics and in silico peptide prediction were applied using a patient-specific data base of SNVs. SYTL4^S363F^ and NCAPG2^P333L^ neoantigens have been detected by MS as previously reported (7) whereas KIF2C^P13L^ has been newly identified through in silico prediction of high affinity HLA binders. Technical improvements of detection sensitivity by MS and data processing are necessary to further enrich for neoantigen identification to fully exploit this technology for standard neoantigen discovery and emphasizes the use of complementary strategies. Experimental affinity measurements of selected predicted peptides showed similar IC_50_ values for all three neoantigens, although no real correlation to predicted affinity was observed for the whole cohort of analyzed peptides. Furthermore, experimental and in silico determined affinity are in our case no accurate immunogenicity predictors. Molecular dynamics confirmed previous reports pointing to the fact that antigen recognition may depend not only on a defined mutation but also on conformational changes affecting the adjacent amino acids (19). Moreover, our data confirm that structural parameters as SASA can be used to improve prediction of suitable neoantigens and may be a valuable addition to comprehensive neoantigen selection algorithms (20).

Interestingly, the somatic mutation coding for KIF2C^P13L^ peptide is in close proximity to another one previously reported. The latter results in an immunogenic decamer which overlaps our neoantigen by 8 amino acids (21). This finding may further support the notion of existing hotspots for peptide presentation and it emphasizes that genes especially activated in cancer, such KIF2C, may preferentially result in peptide ligands (22).

Although several groups have published data describing neoantigen-specific TCRs (23-26), the data shown here provide a much deeper insight. We observed oligoclonal endogenous T-cell responses against two antigens within one patient. We showed recognition of endogenously processed neoantigens as well as tumor reactivity in vitro and in vivo by all identified TCRs. Of particular interest, TCRs with identical specificity showed similar characteristics with regard to temporal and spatial distribution in the body, functional avidity and functionality, suggesting a major role of the antigen driving defined immune responses. Functional avidities of all seven TCRs were in a narrow range, yet, TCRs with specificity for KIF2C^P13L^ showed significant lower avidities compared to the other receptors and a much higher frequency in both metastatic lesions, adjacent lymph nodes as well as the peripheral blood over time. Moreover, despite inferior cytokine secretion compared to SYTL4^S363F^-specific TCRs in vitro, KIF2C^P13L^-specific TCRs demonstrated favorable tumor control in vivo even at low transduction rates. We hypothesize that high-affinity TCRs may be particularly prone to T-cell dysfunction (27,28) as previously demonstrated for CAR-T cells (29). This hypothesis is further corroborated by high PD-L1 expression on patient’s tumor cells within M_Lung_ (7) potentially inducing a PD-L1-mediated dysfunctional state. In vitro data of TCR-transgenic T cells suggest a different activation signature exemplarily tested for SYTL4-TIL1 in comparison to KIF2C-PBC2, with SYTL4-TIL1 displaying a higher PD-1 and an elevated T-bet expression upon neoantigen-specific stimulation. Differences in T-bet upregulation may be associated to skewing CD8+ T cells towards a short-lived effector phenotype and may contribute to a dysfunctional state (30,31). Interestingly, levels of IFN-γ secretion, T-bet and PD-1 expression substantially decreased upon restimulation of SYTL4-TIL1 in contrast to KIF2C-PBC2. These data may indicate a disadvantage of SYTL4-specific T cells with respect to sustained function. Our data therefore suggest that future engineering of TCR and respective transgenic T cells may be considered as a very personalized task for the individual TCR. This may affect defined comprehensive T-cell engineering measures as PD-1 knock-down as well as affinity maturation of tumor-specific TCRs (32-34). Yet, further investigations implementing multi-parametric transcriptional profiling are needed for a deeper understanding and dissection. Advantages of TCRs with lower avidity with respect to persistence have been recently reported for virus-specific T cells (35) and the observations might be explained by maintenance of such TCRs throughout presentation of similar peptides potentially leading to cross-reactive stimulation (36). Interestingly, occurrence of side effects correlates with favorable clinical response in patients treated with immune checkpoint inhibitors, suggesting that cross-reactivity may in fact be to some extent beneficial (37). Despite this, no major side effects associated to Ipilimumab were observed in the clinical history of patient Mel15. In the case of NCAPG2^P333L^ the coding mutation was lost during progression of disease, coinciding with low frequencies of this TCR and loss of detection of this clonotype in M_Lung_. However, despite low frequencies, both SYTL4^S363F^- and NCAPG2^P333L^-specific TCRs could be still isolated from peripheral blood or TILs, indicating the presence of non-terminally exhausted memory T cells with potential for expansion. Moreover, non-malignant lymph nodes may also represent an attractive source for isolation of such neoantigen-specific TCRs.

In retrospect, we have identified seven endogenous neoantigen-specific TCRs with high potential for transgenic modification of T cells, to be used for adoptive T-cell transfer. Our data provide evidence that TCRs display different qualities in vitro and in vivo, which seem to be class or antigen dependent and need to be taken into account in the context of adoptive T-cell transfer of neoantigen-specific TCR-transgenic T cells.

## Acknowledgments

The authors thank patient Mel15 for participating in the study and his continuous support. We also thank A. Stelzl for excellent technical support as well as Jürgen Schlegel and Sandra Baur (TU München, Department of Neuropathology, Munich, Germany) for technical advice.

