## Supplementary Material + Methods, Supplementary Tables, Supplementary Figure Legends for "Spatial and temporal plasticity of neoantigen-specific T-cell responses bases on characteristics associated to antigen and TCR"

**Supplementary Material and Methods**

### Clinical course of Mel15

### Patient Mel15 was diagnosed with malignant melanoma in 2008 and underwent resection of the primary tumor in the same year. In 2013 metastatic disease with pulmonary and intestinal metastases were detected. A biopsy from the lung metastasis (B_Lung_) was performed for histologic confirmation of metastatic status of malignant melanoma. Subsequently, the patient underwent two cycles of chemotherapy resulting in a mixed response with regression of the intestinal metastasis (M_Int_) but progression of lung metastasis (M_Lung_). In November 2013, the patient became eligible for administration of anti-CTLA-4-antibody Ipilimumab for a total of four cycles. In March 2014, restaging revealed response of M_Lung_, whereas M_Int_ progressed and was therefore surgically removed, together with two non-malignant lymph nodes (M_Int_-LN1 and -LN2). The residual M_Lung_ previously targeted by biopsy, initially regressed, but later progressed and was therefore surgically removed in January 2016, together with one adjacent non-malignant lymph node (M_Lung_-LN). Treatment with the PD-1-directed monoclonal antibody Pembrolizumab was subsequently applied for one year and the patient is in complete remission since then (Supplementary Fig. S1).

### Cultivation of cell lines and primary human cells

### All target cell lines were maintained in RPMI 1640, MEM or DMEM (Invitrogen), according to manufacturer's instructions, supplemented with Penicillin and streptomycin antibiotics (Pen/Strep), L-glutamine, non-essential amino acids (NEAA), sodium pyruvate (NaP) and 10% heat-inactivated fetal calf serum (FCS). T cells were cultured in RPMI supplemented with: L-glutamine, 5% human serum and 5% FCS, Hepes, Pen/Strep, NEAA, NaP and gentamycin.

### T-cell antigen-experience recall responses

### For stimulation with peptides, PBMCs were cultured as previously described (1,2). Briefly, frozen PBMCs were thawed and cultured in AIM-V medium supplemented with 100 ng/ml of IL-4 and 100 ng/ml GM-CSF. After 24h, Poly:IC (20 µg/ml), IL-7 (0.5 ng/ml) and peptide pools (1 µM/peptide) were added to the culture. After 20-24h incubation, PBMCs were transferred on a pre-coated IFN-γ ELISpot plate, which was developed after additional 20-24h, prior transfer of PBMCs to culture for expansion of T cells in RPMI medium supplemented with IL-7 (5 ng/ml) and IL-15 (5 ng/ml) for 10-15 days. IFN-γ ELISpot assay was repeated on expanded T cells co-cultured for 72h with T2-A3 cells (E:T = 1:1) pulsed with single peptides from sub-pool A4 shared by Pool 4 and 9.

### Arrangement of peptide pools for immunogenicity assessment of predicted ligands

The first 25 predicted binders to each allele were synthetized and used for stimulation assays, excluding previously identified neoantigens (NCAPG2^P333L^, ranked 24^th^ and SYTL4^S363F^ ranked 6^th^). In total 48 peptides were assembled in 11 pools. Each pool contained 10 peptides: 5 HLA-A03:01-predicted binders (sub-pool A) and 5 HLA-B27:05-predicted binders (sub-pool B) were arranged as depicted in Supplementary Table S3. Sub-pools A5 and B2 contained 9 peptides, due to the exclusion of known immunogenic peptides (1). Pool 11 contained peptide SYTL4^S363F^ and served as positive control for peptide pool approach. Following screening of peptide pools, T cells were expanded for two weeks and co-cultured with single-peptide-pulsed target T2 cells to define the peptides eliciting reactivity in the pool.

**IC_50_ in vitro measurement of predicted peptide candidates and neoantigens**

Binding of selected peptides to HLA was further investigated through a peptide exchange assay. HLA-A03:01, HLA-B27:05, and human β2m (hβ2m) were expressed in E. coli and purified from inclusion bodies as described previously (3). Purified proteins were stored at −80 °C until use. Folding of heavy chain–hβ2m–peptide complexes was performed by diluting purified denatured proteins to a final concentration of 30 µg/mL of heavy chain and 30 µg/mL hβ2m folding buffer (100 mM Tris·Cl, pH 7.5, 0.4 M arginine, 2 mM EDTA, 0.5 mM oxidized glutathione, 5 mM reduced glutathione, and 0.5 mM PMSF) and stirring at 4 °C (2 days for HLA-A03:01, 14 days for HLA-B27:05) with 10 µM high-affinity peptide (KLIETYFJK for HLA-A03:01, and RRKJRRWHL for HLA-B27:05, where J is the photolabile aa residue) (4). After folding, samples were ultracentrifuged at 15,000 × g for 20 min and further purified by gel filtration. For peptide exchange reaction, 0.5 µM HLA class I monomers (HLA–hβ2m– photolabile peptide complex) were exposed to UV radiation (345 nm, 1000 W) for 5 min, followed by incubation for 18 h at 4 °C with 10nM fluorophore-labeled peptide (KLIE-FITC-YFSK for HLA-A3:01 and RRKW-FITC-RWHL for HLA-B27:05, FITC λex = 494 nm, λem = 517 nm) and varying concentration of the measured peptide. The binding was measured by fluorescence anisotropy using a Victor 3V reader (Perkin Elmer) in a black 96-well plate (Corning). Data were fitted, and IC_50_ values were calculated by using GraphPad Prism 7.

**TCR in situ hybridization**

RNA in situ hybridization for identification of neoantigen-specific TCR on FFPE tumor slides was performed using BaseScope^TM^ Detection Reagent Kit v2 - RED (ACD). TCR-specific Probes were designed to anneal on the TCR CDR3 sequence. Probes against PPIB (Cyclophilin B) and a bacterial protein (DapB, Bacillus subtillis) served as respective positive and negative controls. For each paraffin tumor block, slides of 4 µm were trimmed with a microtome. Tissue slides were deparaffinized, pretreated using Protease IV (ACD) and stained with target probe, negative control and positive control probes following the manufacturer’s instructions on an automated immunostainer. For localization of T-cell infiltration, Immunohistochemistry (IHC) was performed using anti-CD3-antibody (clone MRQ, Cell Marque Cat# 103R, RRID AB_2864399) as previously described (1). Slides were scanned on a Leica AT2 (Leica, Wetzlar, Germany) system and visualized using Aperio ImageScope software (v12.4.3).

### HLA-peptide complex structural modelling and SASA calculation

Each model was solvated in a rectangular box of TIP3P (5) water with a minimum distance of 12 Å between solute and box boundary, neutralized, and energy minimized. All modelling steps were performed using AMBER17 (6). The force field FF14SB was selected (7). Subsequently, all 6 models were progressively heated up to 300 K over 3 ns gradually releasing initial positional restraints from the solute (Supplementary Table S4). The last heat-up step was performed in constant number, pressure and temperature (NPT) thermodynamic ensemble whereas the other steps were performed in constant number, volume and temperature (NVT). Temperature was controlled using a Langevin thermostat with a collision frequency (gamma_ln) of 4.00 ps^-1^. Pressure was controlled using a Berendsen barostat (8) with default settings. SHAKE algorithm (9) was applied for all bonds including hydrogen atoms. Particle mesh Ewald (PME) method (10) was used to compute long range electrostatic interactions. A cut-off for non-bonded interactions of 12 Å was applied. After equilibration models were simulated for 200 ns at 300 K using the NPT ensemble with a 1 fs time step. For each model 3 replicas were simulated resulting in a cumulative simulation time of 600 ns per system.

To quantitatively analyse the HLA-p interface, solvent-accessable surface area (SASA) was calculated for each peptide residue and the electrostatic potential of the interaction surface as previously describe (11,12). Briefly, frames were extracted every 200 ps from the last 180 ns of each replicon and analyzed. To select representative structures for visualization, frames were clustered, and central structure of the highest populated cluster was picked. The clustering was based on non-hydrogen peptide atoms using the hierarchical agglomerative algorithm with an epsilon of 2.0 and default settings. Trajectory processing and clustering was conducted with the CPPTRAJ (13) tool provided in AmberTools17 (6). Molecule images were generated using PyMOL (version 1.7.X, Schrödinger, LCC). Electrostatic potential was computed with PyMOL plugin APBS Tools 2.1 (14) using default settings. Hydrogen bonds were identified using the VMD software HBonds plugin (version 1.2) (15).

**In-vitro cytotoxicity assessment**

For monitoring of T-cell mediated killing in vitro, two different adherent cell lines were chosen according to the naturally expressed HLA allotypes. MDST8 cell line expressing HLA-B27:05 was selected for SYTL4^S363F^-specific TCRs; A2058 expressing HLA-A03:01 for KIF2C^P13L^ and NCAPG2^P333L^-specific TCRs. Killing assays were performed with impedance-based xCELLigence assays (ACEA BioSciences) (16). Target cell culturing media was added to each well of 96 well E-Plates (ACEA Biosciences) for background impedance measurement (17). Dissociated adherent target cells were seeded on E-Plate at different densities depending on the cell line (A2058 – 50,000/well; MDST8 – 20,000/well). Cell density on plates is measured within the RTCA MP instrument inside a cell culture incubator for 24h. For addition of T cells, data acquisition was paused and effector cells were added. Data recording was initiated immediately at 15-min intervals for the first 8 h and then at 30-min intervals for remaining 16 h. Target cells only as well as target cells and non-transduced T cells served as controls. For calculation of percentage of cytolysis, the following formula was adopted:
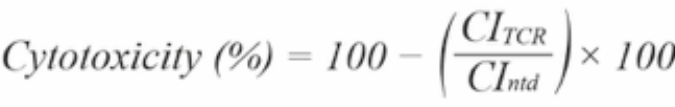
.

**Activation/dysfunction patterns of stimulated and restimulated enriched TCR-transgenic T cells**

A near-infrared fluorescent protein (iRFP) linked with a T2A element was cloned downstream of TCR constructs KIF2-PBC2 and SYTL4-TIL1 to allow for flow-cytometry based enrichment of TCR-transduced T cells. Retroviral transduction was performed as stated in the method section and iRFP-positive T cells were sorted on a FACSAria III Cell Sorter (BD Biosciences). Sorted T cells were coincubated with minigene-transduced U-698-M in an E:T ratio of 1:1. To analyze repeated antigen exposure, transduced and enriched T cells were cocultured with irradiated U-698-M expressing mutated tandem minigene (E:T = 1:1), fed with IL-7 and IL-15 (5 ng/ml each) every three days and restimulated with respective target cells after 11 days. For flow cytometry assessment, T cells were stained with anti-CD8, TCRmu and Zombie UV Fixable Viability Kit (BioLegend). Expression of activation/dysfunction markers was assessed using anti-PD-1-antibody (BioLegend, Cat# 329930, RRID AB_2563443) and, in combination with True-Nuclear Transcription Factor buffer set (BioLegend), anti-T-bet (BioLegend, Cat# 644816, RRID AB_10959653). Proliferation after 6 days of target stimulation was assessed with CellTrace Violet Cell Proliferation Kit, (ThermoFisher) according to the manufacturer’s recommendations. Flow cytometry data acquisition and analysis was performed as stated in respective section within the main manuscript.

**Supplementary Figure Legends**

**Figure S1.**

Clinical course of disease of patient Mel15. Patient Mel15 was diagnosed with malignant melanoma in 2008 and underwent resection of the primary tumor that year. In 2013, pulmonary and intestinal metastases were detected and a lung biopsy (BLung) was performed for histological analysis. Carboplatin and paclitaxel were administered, resulting in a mixed response of metastatic disease. The patient was subsequently treated with the anti-CTLA-4-antibody Ipilimumab. In 2014, the progressing abdominal lesion (MInt) and two non-malignant lymph nodes (LNs) were surgically removed. In 2016, one primarily responding lung metastasis (MLung) progressed and was resected with one adjacent non-malignant LN. Treatment with Pembrolizumab was subsequently administered for 18 months. The patient is currently in complete remission without therapy. Analyses performed on tissue samples and time points of blood withdrawals are depicted.

**Figure S2.**

Detection of immunogenicity of predicted neoantigen KIF2CP13L in autologous PBMCs of patient Mel15. **A**, PBMCs derived from time point 925 were cultured in presence of IL-4 and GM-CSF following addition of Poly:IC, IL-7 and peptide pools (1 µM/peptide). After 24h, IFN-γ ELISpot assay was performed. **B**, IFN-γ ELISpot assay with T cells expanded after stimulation with Pool 4 and 9 (Supplemental Table 3). Expanded T cells were incubated with T2-A3 cells (E:T = 1:1) pulsed with single peptides of subpool A4 shared by Pool 4 and 9 for 72h. Subpool B4 containing peptides only present in Pool 4 but not Pool 9, served as negative control.

**Figure S3.**

Mutations result in distinct antigen features as shown by structural modeling. **A**, **B** and **C**, residue specific SASA values of the neoantigen (red) and its respective WT (blue) are plotted on the left side. Corresponding superimposed representative structures as extracted from MD simulations are shown on the right side for SYTL4^S363F^(**A**), KIF2C^P13L^(**B**), and NCAPG2^P333L^(**C**). HLA peptide complexes (right side) are shown in surface representation with HLA (light grey), peptide (dark grey) and central positions of increased SASA in red (neoantigen) and blue (WT). Latter positions are emphasized in the SASA plots by black arrows and dashed line ellipses. **A** and **B**, mutated residues are depicted in sticks and transparent spheres to illustrate physicochemical differences. In **C**, the buried mutation site is indicated by the arrow. SASA mean value and SD of each individual peptide aa was calculated out of three replicates. * p < 0.05; ** p < 0.01.

**Figure S4.**

Expression of transgenic neoantigen-specific TCRs on the surface of human CD8+ T cells assessed by TCRmu staining. Representative flow cytometry plots of single-cell suspension of TCR-transduced-CD8+ T cells stained (7-10 days after transduction) for CD8, TCRmu, and 7-AAD; the dot plots were gated on living cells (7-AAD-) and CD8-TCRmu+ T cells. The numbers in the gate represent the percentage of CD8+-T cells expressing the murinized TCR. Gates were set on the basis of isotype control for TCRmu-Ab, for each single TCR. Transduction efficiency was assessed for all transgenic T cells showing consistent results for the different TCR constructs.

**Figure S5.**

Expression of transgenic neoantigen-specific TCRs on the surface of CD8^+^ T cells assessed by multimer staining. Representative flow cytometry plots show stainings with indicated multimers: HLA-B27:05-SYTL4 (**A**), HLA-A03:01-KIF2C (**B**) and HLA-A03:01-NCAPG2 (**C**). TCR-transduced-CD8^+^ T cells were then stained for CD8 and 7-AAD; the dot plots were gated on living cells (7-AAD^-^). The values in the gate represent the percentage of multimer-positive CD8^+^-T cells. The quadrants were set on the basis of non-transduced T cells and isotype control. One representative out of three experiments with T cells from two different healthy donors is shown.

**Figure S6.**

Functional avidity of neoantigen-specific TCRs. Representative plots of IFN-γ secretion of TCR-transduced T cells upon stimulation with T2 target cells pulsed with titrated peptide concentrations of SYTL4^S363F^ (**A**), KIF2C^P13L^ (**B**) and NCAPG2^P333L^ (**C**). EC_50_ values of all experiments were calculated by fitting a non-linear curve and results are summarized in Fig. 1D.

**Figure S7.**

TCR-transduced CD8^+^ T cells mediate specific lysis of target cells expressing defined neoantigens. **A**, **B** and **C**, growth of target cells transduced with MUT/WT minigenes is monitored for ca. 24h (impedance measurement every 15 min) before addition of transgenic T cells to the culture and observation of cytolysis. MDST8MUT/WT cells were seeded at density 20,000/well and 40,000 TCR-T cells/well were added at time 0 (**A**). A2058MUT/WT cells were seeded at density 50,000/well and 100,000 TCR-T cells/well were added (**B** and **C**). Cell index values of target cell lines are depicted on left y axis. Percentage of target cell lysis, calculated on non-transduced T cell, is depicted on right y axis and indicated by the colored line. **D**, Comparison of cytolysis mediated by all TCRs. TCR transduction efficiency is indicated in the figure legend.

**Figure S8.**

Investigation of peptide-dependent and – independent HLA alloreactivity of neoantigen-specific TCR. Reactivity of all defined TCRs was tested to different LCLs expressing common HLA allotypes, pulsed with 1 μM mutated peptides SYTL4^S363F^, KIF2C^P13L^ and NCAPG2^P333L^ (pLCL) in comparison to non-pulsed condition (LCL). All experiments were performed three times on two different healthy-donor-derived sets of transduced T cells showing consistent results. For cocultures, E:T ratio was 1:1 and read-out consisted of IFN-γ secretion, assessed by ELISA on culture supernatants after 20h incubation

**Figure S9.**

Venn diagrams with distribution of TCR CDR3 sequences. **A**, Overlap of TCR CDR3 clonotypes obtained from TCR-β sequencing on B_Lung_, M_Int_ and M_Lung_. **B**, Overlap of TCR CDR3 clonotypes from removed metastases and associated lymph nodes. Arrows indicate shared compartments in which defined TCRs with validated neoantigen-specificity were detectable. The overlap was calculated using the Venn diagram tool from Bioinformatics & Systems Biology available online (http://bioinformatics.psb.ugent.be/webtools/ Venn/).

**Figure S10.**

RNA in situ hybridization of neoantigen-specific T cells within tumor tissue MLung. **A**, CDR3-specific probe visualizes T cells expressing TCR KIF2C-PBC2, which show polarization of detected RNA within single cells. **B**, anti-CD3 staining show an inhomogenous T-cell infiltrate. **C**, Negative control probe DapB (Bacillus subtillis). **D**, positive control probe Cyclophilin B.

**Figure S11.**

Expanded TIL from M_Lung_ of Mel15 (1) were analyzed for specific responses against mutated peptide ligands. IFN-γ secretion upon stimulation with peptide-pulsed autologous LCL line and analyzed by IFN-γ ELISpot.

**Figure S12.**

Longitudinal monitoring of activation of TCR-transgenic T cells upon neoantigen encountering. **A-B**, percentage of T-bet positive cells within TCRmu^+^CD8^+^ fraction of TCR-transduced T cells after 24 and 96 hours of coculture with U-698-M expressing the mutated (**A**) or wildtype (**B**) tandem minigene. **C-D**, T-bet expression of SYLT4-TIL1-iRFP and KIF2C-PBC2-iRFP transduced T cells as shown in Fig. 5C including analysis after 96 hours after coincubation with mutated (**C**) or wildtype (**D**) transduced U-698-M.

**Supplementary Tables**

| Rank | Gene | Peptide seq (aa) | Mutation (aa) | Predicted affinity (nM) | Experimental IC_50_ (nM) |
| --- | --- | --- | --- | --- | --- |
| **HLA-A03:01** | | | | | |
| 1 | *CRMP1* | KIFNFYPRK | L > F | 6.8 | 11.5 |
| 2 | *FUT9* | KMKNFFFTK | S > F | 7.4 | 135.7 |
| 3 | *ANO7* | RMLRRRAQK | E > K | 8.7 | 791.3 |
| 4 | *ADAMDEC1* | TLYSPRGEK | E > K | 9.2 | 416.9 |
| 5 | *ASAH2* | AMYQRAKLK | S > L | 9.5 | 27.1 |
| 6 | *SLC5A3* | SLLTPPSTK | P > S | 9.6 | 211.4 |
| 7 | *CRYBB1* | RLMFFRPIK | S > F | 9.8 | 57.6 |
| 8 | *ZNF234* | SLYLKIHLK | L > K | 11 | -- |
| 9 | *ZNF234* | KIYAAGTFY | H > Y | 11.2 | 975 |
| 10 | *DNAH5* | YLFFIQGYK | S > F | 12.5 | 932.8 |
| 11 | *TENM1* | TTYSPIGEK | G > E | 14.5 | 81.51 |
| 12 | *NCAPG2* | RLYKLILWR | P > L | 14.7 | 707.1 |
| 13 | *TGM5* | KTYPCKIFY | S > F | 16.6 | 248.1 |
| 14 | *FHDC1* | SLQPRGSFK | P > S | 18.2 | 36.0 |
| 15 | *CCDC7* | KVINLSPFK | E > K | 18.6 | 474.3 |
| 16 | *PCDHGA10* | CLFFGIPWK | S > I | 19.2 | 2391 |
| 17 | *SGK3* | KQFSAMALK | P > S | 21.4 | 208.2 |
| 18 | *KIF2C* | RLFLGLAIK | P > L | 21.6 | 277.4 |
| 19 | *AKAP6* | KLKLPIIMK | M > I | 23.3 | 35.1 |
| 20 | *FREM1* | LLINRGFSK | D > N | 25.2 | 931.2 |
| 21 | *UVSSA* | RLKCPFYGK | H > Y | 26.1 | 249 |
| 22 | *USP37* | KVMTDPSRK | A > V | 28.7 | 16,29 |
| 23 | *ARHGEF17* | RIAGKALKK | P > L | 31.5 | 669 |
| 24 | *NCAPG2* | KLILWRGLK | P > L | 32.6 | 263.9 |
| 25 | *ZFN226* | KLYQCNECK | S > L | 32.7 | 3502 |

**Table S1.** Measured binding activity of top 25 neoantigen candidates for HLA-A03:01 predicted by NetMHC4.0.

| Rank | Gene | Peptide seq (aa) | Mutation (aa) | Predicted affinity (nM) | Experimental IC_50_ (nM) |
| --- | --- | --- | --- | --- | --- |
| **HLA-B27:05** | | | | | |
| 1 | *TECTB* | R**R**FSSLYSF | G > R | 11.5 | 48.0 |
| 2 | *CLDN24* | RRLLILG**R**I | G > R | 11.5 | 311.0 |
| 3 | *KAT2A* | FRMF**L**TQGF | P > L | 15 | 2.0 |
| 4 | *TMPRSS11E* | ARWTA**F**FGV | S > F | 17.1 | 525.0 |
| 5 | *SDK2* | GRWALHSA**F** | S > F | 17.5 | 68.0 |
| 6 | *SYTL4* | GRIAF**F**LKY | S > F | 18.4 | 224.0 |
| 7 | *Ac087289.2, MRPL38, Ac087289.3^a^* | KRF**L**HRQPL | P > L | 20.2 | 73.0 |
| 8 | *B3GAT1* | A**R**FAVNLRL | G > R | 20.7 | 8.0 |
| 9 | *ITLN1* | WRNS**F**LLRY | S > F | 24 | 29.0 |
| 10 | *Af241726.2, CYBB^a^* | YR**I**YDIPPK | V > I | 24 | 60.0 |
| 11 | *KIF2C* | ARLF**L**GLAI | P > L | 25.7 | -- |
| 12 | *P2RX1* | YRHLFKVF**R** | G > R | 26.6 | 70.0 |
| 13 | *SCN2A* | FRFFTR**K**SL | E > K | 26.9 | 4.0 |
| 14 | *RNF216* | RRHCRSY**N**R | D > N | 27.6 | 55.0 |
| 15 | *CLDN24* | KRRLLILG**R** | G > R | 28.0 | 111.0 |
| 16 | *ZFN234* | FRQSLYL**K**I | L > K | 29.2 | 7.0 |
| 17 | *NEK10* | RRTQRYFM**K** | E > K | 29.3 | 46.0 |
| 18 | *ERCC5* | FRI**C**PIFVF | R > C | 32.3 | 877.0 |
| 19 | *DSG1* | **K**RTNVGILK | E > K | 33.3 | 136.0 |
| 20 | *LIPH* | LRILR**I**KLR | M > I | 35.6 | 109.0 |
| 21 | *SI* | KR**H**EVPVPL | Y > H | 36.8 | 907.0 |
| 22 | *MBOAT1* | HRY**F**FFVAM | S > F | 37.5 | 353.0 |
| 23 | *SCN4A* | FRF**F**ATPAL | S > F | 38.3 | 13.0 |
| 24 | *INO80C* | LRFS**I**IEEF | T > I | 45.9 | -- |
| 25 | *STON2* | SRVILF**S**PL | N > S | 46.0 | -- |
| a) Overlapping genes | | | | | |

**Table S2.** Measured binding activity of top 25 neoantigen candidates for HLA-B27:05 predicted by NetMHC4.0.

| **Subpool A** | **A03:01 binders** | **Subpool B** | **B27:05 binders** | **Pools** |
| --- | --- | --- | --- | --- |
| A1 | KIFN**F**YPRK | B1 | R**R**FSSLYSF | Pool 1 |
|  | KMKNFF**F**TK |  | RRLLILG**R**I |  |
|  | RMLRRRAQ**K** |  | FRMF**L**TQGF |  |
|  | TLYSPRGE**K** |  | ARWTA**F**FGV |  |
|  | AMYQRAK**L**K |  | GRWALHSA**F** |  |
| A2 | SLLTPP**S**TK | B2 | - | Pool 2 |
|  | RLM**F**FRPIK |  | KRF**L**HRQPL |  |
|  | SLYL**K**IHLK |  | A**R**FAVNLRL |  |
|  | KI**Y**AAGTFY |  | WRNS**F**LLRY |  |
|  | YL**F**FIQGYK |  | YR**I**YDIPPK |  |
| A3 | TTYSPIG**E**K | B3 | ARLF**L**GLAI | Pool 3 |
|  | RLYK**L**ILWR |  | YRHLFKVF**R** |  |
|  | KTYPCKI**F**Y |  | FRFFTR**K**SL |  |
|  | **S**LQPRGSFK |  | RRHCRSY**N**R |  |
|  | KVINLSPF**K** |  | KRRLLILG**R** |  |
| A4 | CLFFG**I**PWK | B4 | FRQSLYL**K**I | Pool 4 |
|  | KQF**S**AMALK |  | RRTQRYFM**K** |  |
|  | RLF**L**GLAIK |  | FRI**C**PIFVF |  |
|  | KLKLP**I**IMK |  | **K**RTNVGILK |  |
|  | LLI**N**RGFSK |  | LRILR**I**KLR |  |
| A5 | RLKCPF**Y**GK | B5 | KR**H**EVPVPL | Pool 5 |
|  | K**V**MTDPSRK |  | HRY**F**FFVAM |  |
|  | RIAGKA**L**KK |  | FRF**F**ATPAL |  |
|  | - |  | LRFS**I**IEEF |  |
|  | K**L**YQCNECK |  | SRVILF**S**PL |  |
| A1 | KIFN**F**YPRK | B2 | - | Pool 6 |
|  | KMKNFF**F**TK |  | KRF**L**HRQPL |  |
|  | RMLRRRAQ**K** |  | A**R**FAVNLRL |  |
|  | TLYSPRGE**K** |  | WRNS**F**LLRY |  |
|  | AMYQRAK**L**K |  | YR**I**YDIPPK |  |
| A2 | SLLTPP**S**TK | B3 | ARLF**L**GLAI | Pool 7 |
|  | RLM**F**FRPIK |  | YRHLFKVF**R** |  |
|  | SLYL**K**IHLK |  | FRFFTR**K**SL |  |
|  | KI**Y**AAGTFY |  | RRHCRSY**N**R |  |
|  | YL**F**FIQGYK |  | KRRLLILG**R** |  |
| A3 | TTYSPIG**E**K | B4 | FRQSLYL**K**I | Pool 8 |
|  | RLYK**L**ILWR |  | RRTQRYFM**K** |  |
|  | KTYPCKI**F**Y |  | FRI**C**PIFVF |  |
|  | **S**LQPRGSFK |  | **K**RTNVGILK |  |
|  | KVINLSPF**K** |  | LRILR**I**KLR |  |
| A4 | CLFFG**I**PWK | B5 | KR**H**EVPVPL | Pool 9 |
|  | KQF**S**AMALK |  | HRY**F**FFVAM |  |
|  | RLF**L**GLAIK |  | FRF**F**ATPAL |  |
|  | KLKLP**I**IMK |  | LRFS**I**IEEF |  |
|  | LLI**N**RGFSK |  | SRVILF**S**PL |  |
| A5 | RLKCPF**Y**GK | B1 | R**R**FSSLYSF | Pool 10 |
|  | K**V**MTDPSRK |  | RRLLILG**R**I |  |
|  | RIAGKA**L**KK |  | FRMF**L**TQGF |  |
|  | - |  | ARWTA**F**FGV |  |
|  | K**L**YQCNECK |  | GRWALHSA**F** |  |
| A1 | SLLTPP**S**TK | B2 | GRIAF**F**LKY | Pool 11 |
|  | RLM**F**FRPIK |  | KRF**L**HRQPL |  |
|  | SLYL**K**IHLK |  | A**R**FAVNLRL |  |
|  | KI**Y**AAGTFY |  | WRNS**F**LLRY |  |
|  | YL**F**FIQGYK |  | YR**I**YDIPPK |  |

**Table S3**. Predicted peptide pool arrangement for immunogenicity assessment.

| **Step** | **Time (ps)** | **Restraint atoms** | **Restraint force constant**  **(kcal mol-1 Å-2)** | **Initial temperature (K)** | **Target temperature (K)** | **Thermodynamic ensemble^d^** | **Barostat** |
| --- | --- | --- | --- | --- | --- | --- | --- |
| 1 | 10 | all^a^ | 2.39 | 0 | 0 | NVT | - |
| 2 | 50 | all^a^ | 2.39 | 2.5 | 5 | NVT | - |
| 3 | 50 | all^a^ | 2.39 | 5 | 10 | NVT | - |
| 4 | 50 | all^a^ | 2.39 | 10 | 20 | NVT | - |
| 5 | 50 | bb^b^ | 2.39 | 25 | 50 | NVT | - |
| 6 | 100 | bb^b^ | 2.39 | 50 | 100 | NVT | - |
| 7 | 100 | bb^b^ | 2.39 | 100 | 200 | NVT | - |
| 8 | 100 | bb^b^ | 0.24 | 100 | 200 | NVT | - |
| 9 | 200 | mh-bb^c^ | 0.24 | 100 | 200 | NVT | - |
| 10 | 200 | mh-bb^c^ | 0.24 | 150 | 300 | NVT | - |
| 11 | 590 | - | - | 150 | 300 | NVT | - |
| 12 | 1500 | - | - | 150 | 300 | NPT | Berendsen |
| a) All MHC and peptide atoms b) Heavy (C,CA,O,N) backbone (bb) atoms of MHC and peptide c) Heavy backbone atoms of MHC (mh-bb) d) NVT: constant number, volume and temperature; NPT: constant number, pressure and temperature | | | | | | | |

**Table S4**. Heat-up parameters used for MD simulations.

| **Name cell line** | **Alias** | **HLA-A** | **HLA-B** | **HLA-C** |
| --- | --- | --- | --- | --- |
| HOM2 | LCL 01 | 03:01 | 27:05:00 | 01:02 |
| SWEIG007 | LCL 02 | 29:02:00 | 40:02:00 | 02:02 |
| AMALA | LCL 03 | 02:17 | 15:01 | 03:03 |
| OZB | LCL 04 | 02:09/03:01 | 35:01/38:01 | 04:01/12:03 |
| RSH | LCL 05 | 68:02/30:01 | 42:01:00 | 17:01 |
| KLO | LCL 06 | 02:08/01:01 | 50:01/08:01 | 07:01/06:02 |
| LWAGS | LCL 07 | 33:01:00 | 14:02 | 08:02 |
| - | LCL 08 | 02:01 | 07:02/15:01 | 30:4/12:03 |
| BM21 | LCL 09 | 01:01 | 41:01:00 | 17:01 |

**Table S5**. HLA-allotypes of lymphoblastoid cell lines

| **TRBD** | 2*01 | 2*01 | 2*01 | 1*01 | 2*01F | 1*01F | 1*01 |
| --- | --- | --- | --- | --- | --- | --- | --- |
| **TRBJ** | 2-5*01 | 2-3*01 | 2-5*01 | 2-3*01 | 2-1*01F | 2-3*01F | 1-6*02 |
| **TRBV** | 7-8*01 | 27*01 | 6-2*01 | 12-3*01 | 7-6*01F | 10-3*02F | 15*02 |
| **TRAJ** | 42*01 | 52*01 | 34*01 | 43*01 | 31*01F | 6*01F | 30*01 |
| **TRAV** | 38-2/DV8*01 | 35*02 | 9-2*02 | 8-3*01 | 14/DV4*02F | 12-2*01F | 12-2*02 |
| **TCR** | SYTL4-TIL1 | SYTL4-TIL2 | SYTL4-PBC1 | SYTL4-PBC2 | KIF2C-PBC1 | KIFC-PBC2 | NCAPG2-PBC1 |
| **Source** | TILs | | PBMCs | | | | |
| **Specificity** | SYTL4^S363F^ | | | | KIF2C^P13L^ | | NCAPG2^P333L^ |

**Table S6**. Characteristics of neoantigen-specific TCRs.

| **TCR** | **Recognition motif^a^** | **Number of antigens^b,c^** |
| --- | --- | --- |
| SYTL4-TIL1 | X-R-I-A-F-F-X-X-X | 6 |
| SYTL4-TIL2 | X-R-I-A-F-F-X-X-X | 6 |
| SYTL4-PBC1 | X-R-I-A-F-F-X-X-X | 6 |
| SYTL4-PBC2 | X-R-I-A-F-F-X-X-X | 6 |
| KIF2C-PBC1 | X-X-X-L-X-L-A-I-K | 60 |
| KIF2C-PBC2 | X-X-X-L-X-L-X-I-K | > 400 |
| NCAPG2-PBC1 | K-X-X-L-W-R-X-X-K | 4 |
| a) Recognition motifs are defined through T cell IFN-γ production in response to alanine/threonine scanned cognate epitopes  b) Number of human proteins containing matching recognition motif according to ScanProSite  c) Results derived from protein sequence database UniprotKB, Swiss-Prot (splice variants included) | | |

**Table S7.** Number of human proteins potentially identified by neoantigen-specific TCR according to their recognition motif.

| **TCR** | **M_Int_ (%)** | **M_Int_-LN1 (%)** | **M_Int_-LN2 (%)** | **M_Lung_ (%)** | **M_Lung_-LN (%)** |
| --- | --- | --- | --- | --- | --- |
| SYTL4-TIL1 | 0.041 | 0.036 | 0.012 | 0.039 | 0.005 |
| SYTL4-TIL2 | 0.079 | 0.022 | 0.019 | 0.017 | 0.026 |
| SYTL4-PBC1 | 0.152 | - | 0.019 | 0.024 | 0.034 |
| SYTL4-PBC2 | 0.029 | 0.014 | - | 0.020 | 0.021 |
| KIF2C-PBC1 | 0.560 | 0.029 | 0.037 | 0.629 | 0.023 |
| KIF2C-PBC2 | 0.115 | 0.007 | 0.019 | 0.800 | 0.070 |
| NCAPG2-PBC1 | 0.003 | 0.007 | - | - | - |

**Table S8.** Productive frequency (%) of neoantigen-specific clonotypes in tumor metastases and lymph nodes detected by TCR-β sequencing.

| **TCR** | **d142 (%)** | **d546 (%)** | **d796 (%)** | **d945 (%)** | **d1120 (%)** | **d1519 (%)** |
| --- | --- | --- | --- | --- | --- | --- |
| SYTL4-TIL1 | 0.0018 | 0.0016 | 0.0022 | 0.0020 | 0.0008 | 0.0010 |
| SYTL4-TIL2 | - | - | 0.0015 | 0.0007 | 0.0008 | 0.0010 |
| SYTL4-PBC1 | 0.0053 | 0.0132 | 0.0112 | 0.0099 | 0.0115 | 0.0094 |
| SYTL4-PBC2 | - | 0.0008 | 0.0015 | 0.0007 | 0.0008 | - |
| KIF2C-PBC1 | 0.1489 | 0.1672 | 0.3273 | 0.5449 | 0.4715 | 0.2505 |
| KIF2C-PBC2 | 0.0132 | 0.0311 | 0.0509 | 0.0365 | 0.0376 | 0.0136 |
| NCAPG2-PBC1 | 0.0247 | 0.0156 | 0.0187 | 0.0205 | 0.0230 | 0.0167 |

**Table S9.** Productive frequency (%) of neoantigen-specific clonotypes detected by TCR-β sequencing on peripheral blood collected at different time points indicated as days (d) after start of Ipilimumab treatment.
