## Supplementary Figures 1-12 for "Spatial and temporal plasticity of neoantigen-specific T-cell responses bases on characteristics associated to antigen and TCR"

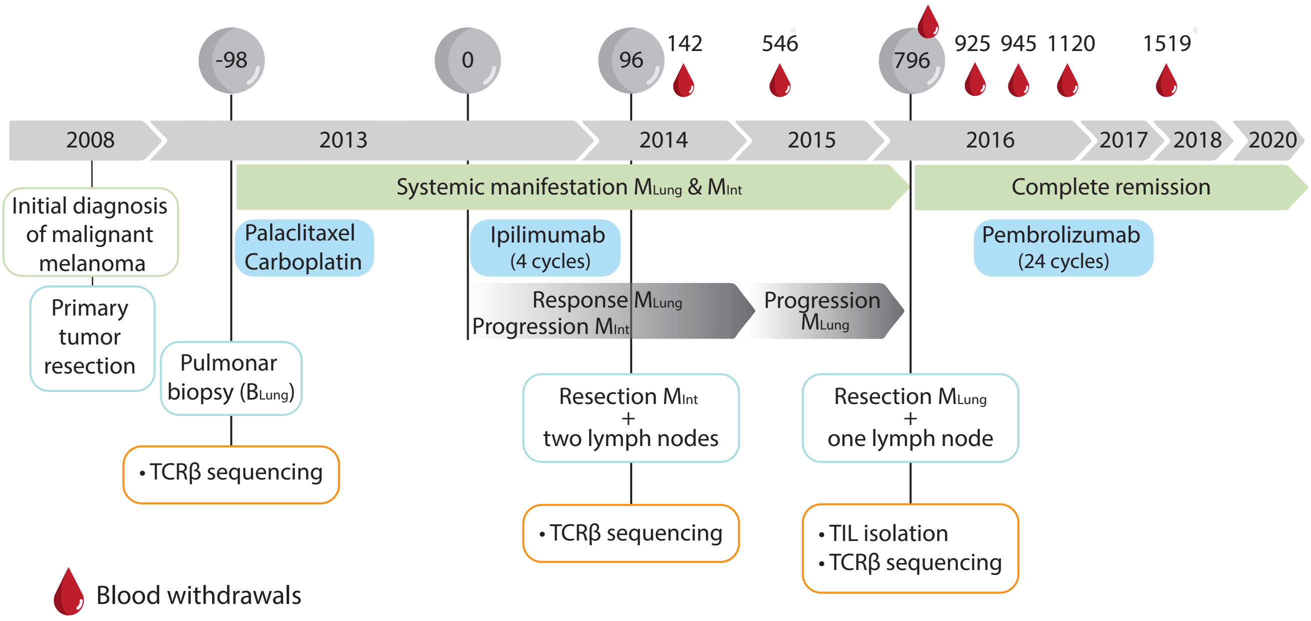
-98
0
96
142
546
796
925
945
1120
1519
2008
2013
2014
2015
2016
2017
2018
2020
Initial diagnosis of malignant melanoma
Primary tumor resection
Systemic manifestation MLung & MInt
Complete remission
Palaclitaxel Carboplatin
Ipilimumab (4 cycles)
Pembrolizumab (24 cycles)
Response MLung Progression MInt
Progression MLung
Pulmonar biopsy (BLung)
Resection MInt + two lymph nodes
Resection MLung + one lymph node
• TCRβ sequencing
• TCRβ sequencing
• TIL isolation
• TCRβ sequencing
Blood withdrawals

Figure S1

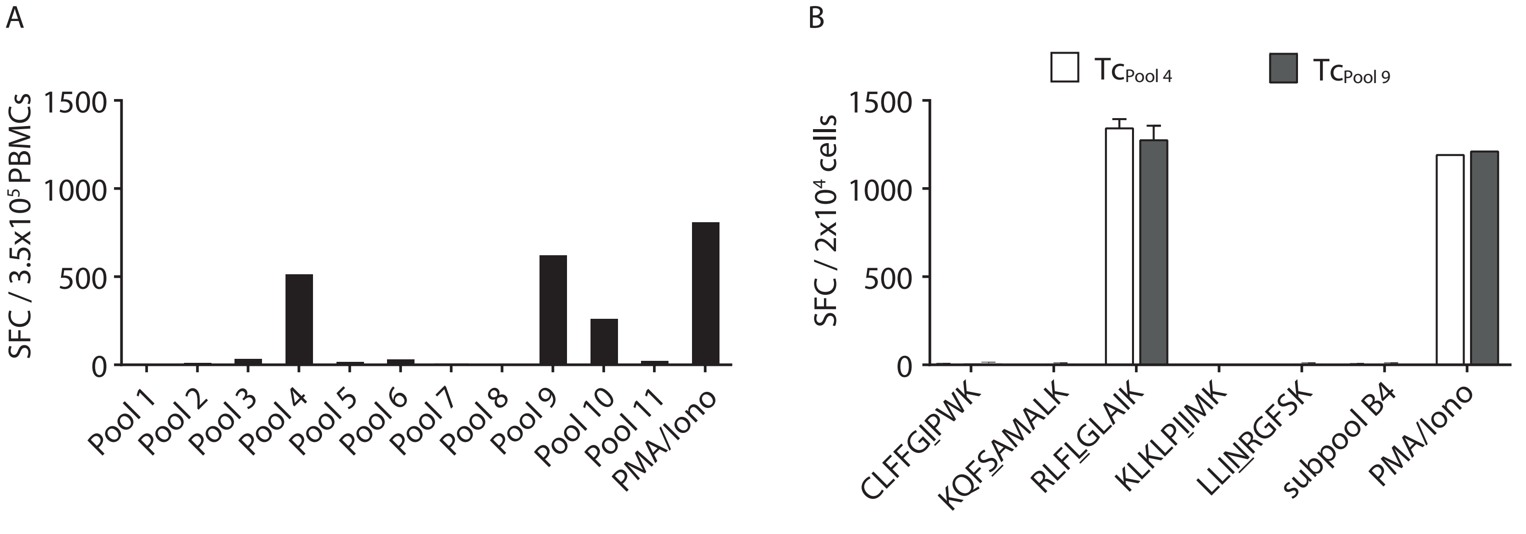
A
SFC / 3.5x10^5 PBMCs
1500
1000
500
0
Pool 1
Pool 2
Pool 3
Pool 4
Pool 5
Pool 6
Pool 7
Pool 8
Pool 9
Pool 10
Pool 11
PMA/Iono
B
Tc Pool 4
Tc Pool 9
SFC / 2x10^4 cells
1500
1000
500
0
CLFFGIPWK
KQFSAMALK
RLFLGLAIK
KLKLPIIMK
LLINRGFSK
subpool B4
PMA/Iono

Figure S2

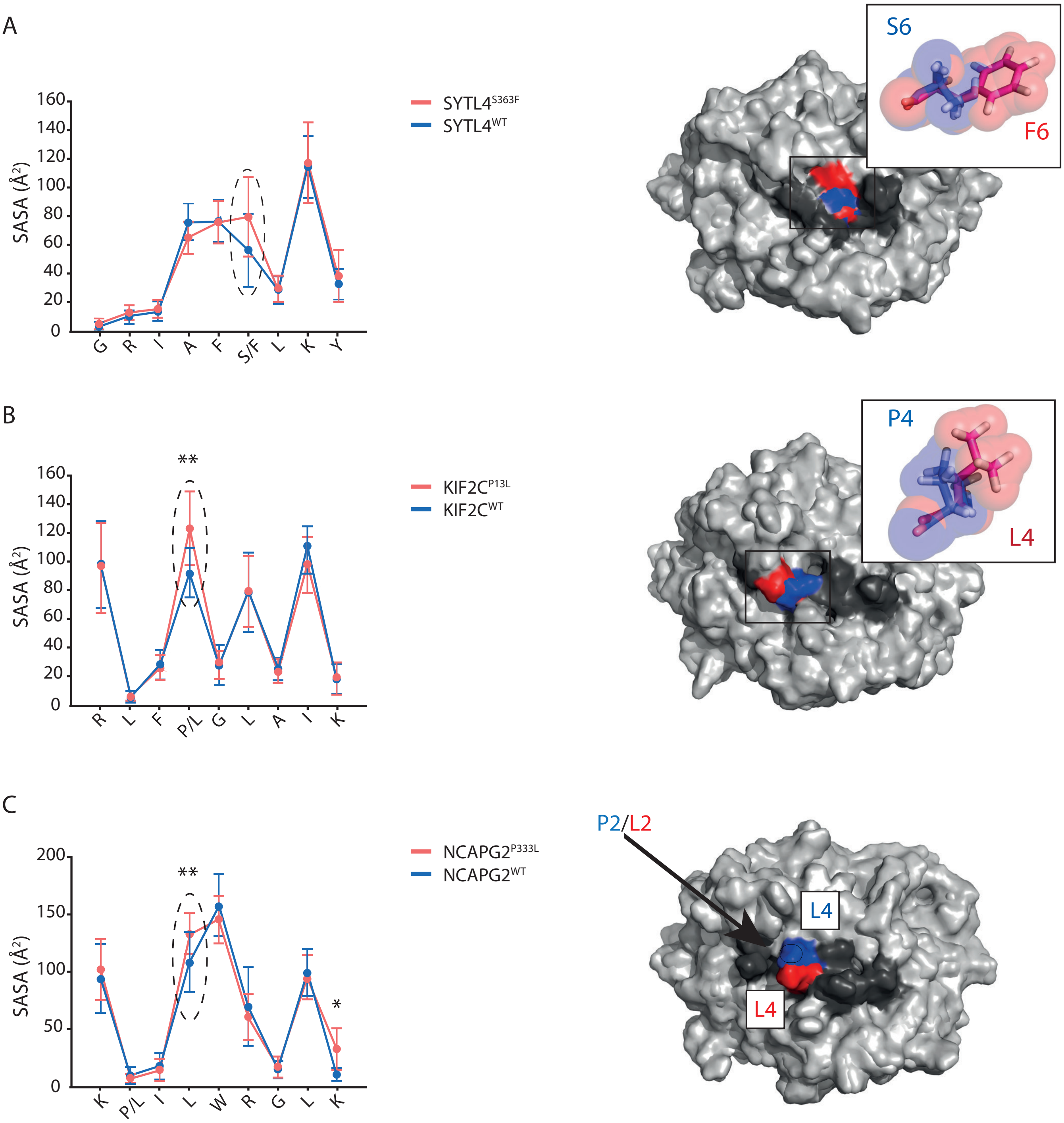
A
SYTL4S363F
SYTL4WT
SASA (Å²)
160
140
120
100
80
60
40
20
0
G
R
I
A
F
S/F
L
K
Y
S6
F6
B
**
KIF2CP13L
KIF2CWT
SASA (Å²)
R
L
F
P/L
G
L
A
I
K
P4
L4
C
**
*
NCAPG2P333L
NCAPG2WT
SASA (Å²)
200
150
100
50
0
K
P/L
I
L
W
R
G
L
K
P2/L2
L4
L4

Figure S3

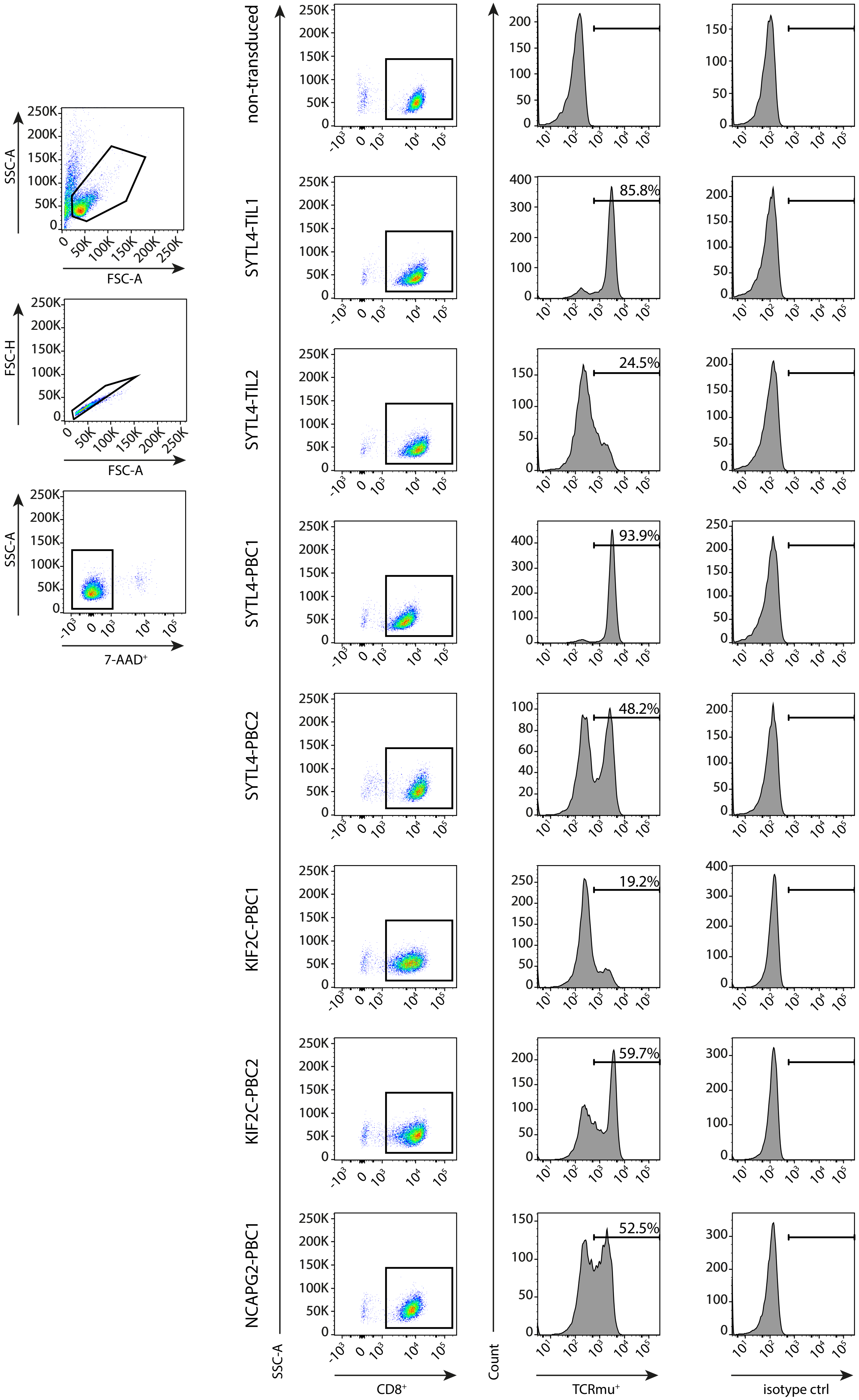

SSC-A
FSC-A
FSC-H
FSC-A
SSC-A
7-AAD+
non-transduced
SYTL4-TIL1
SYTL4-TIL2
SYTL4-PBC1
SYTL4-PBC2
KIF2C-PBC1
KIF2C-PBC2
NCAPG2-PBC1
85.8%
24.5%
93.9%
48.2%
19.2%
59.7%
52.5%
SSC-A
CD8+
Count
TCRmu+
isotype ctrl


Figure S4

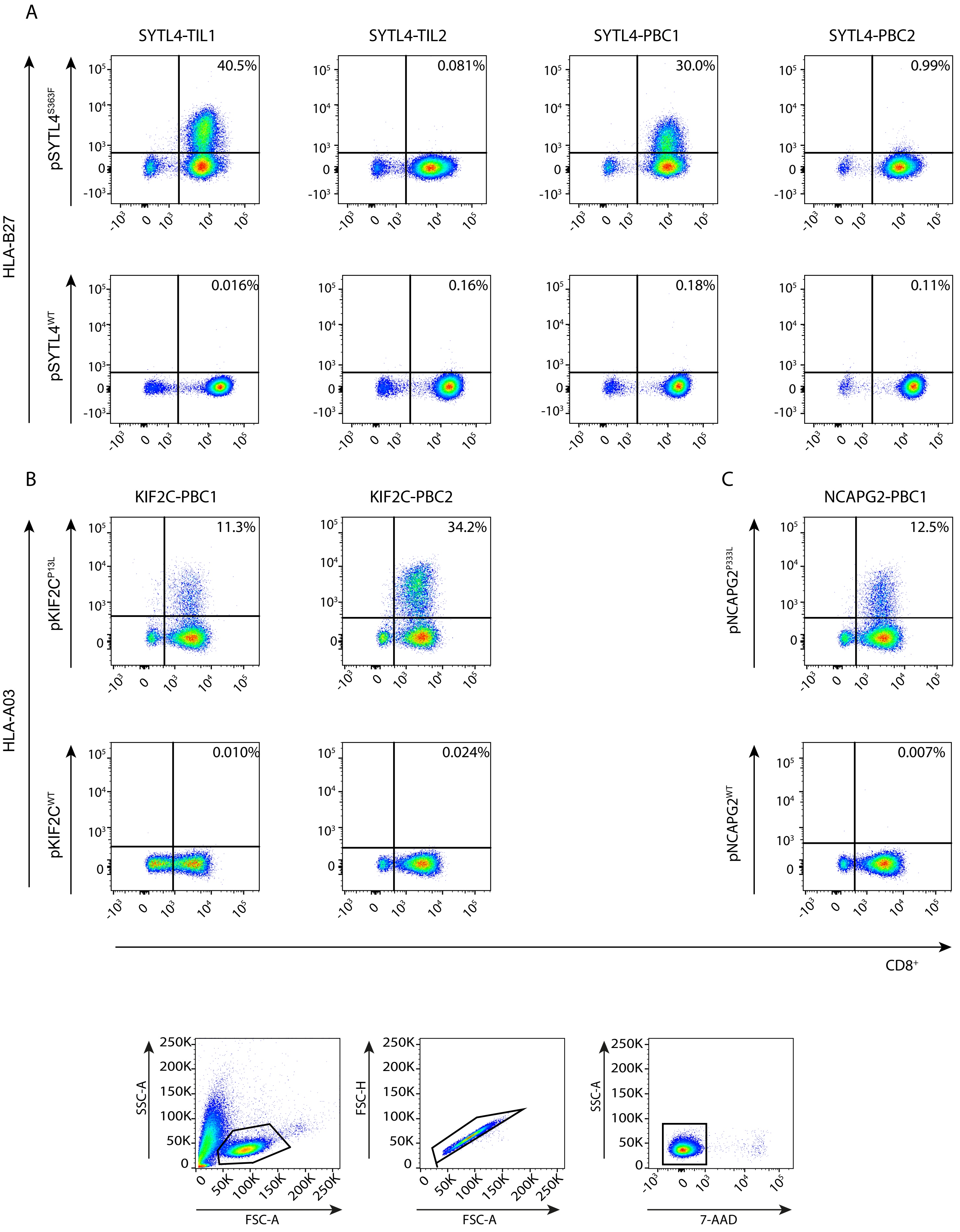
A
SYTL4-TIL1
SYTL4-TIL2
SYTL4-PBC1
SYTL4-PBC2
HLA-B27
pSYTL4S363F
pSYTL4WT
40.5%
0.081%
30.0%
0.99%
0.016%
0.16%
0.18%
0.11%
B
KIF2C-PBC1
KIF2C-PBC2
HLA-A03
pKIF2CP13L
pKIF2CWT
11.3%
34.2%
0.010%
0.024%
C
NCAPG2-PBC1
pNCAPG2P333L
pNCAPG2WT
12.5%
0.007%
CD8+
SSC-A
FSC-A
FSC-H
7-AAD
250K
200K
150K
100K
50K
0

Figure S5

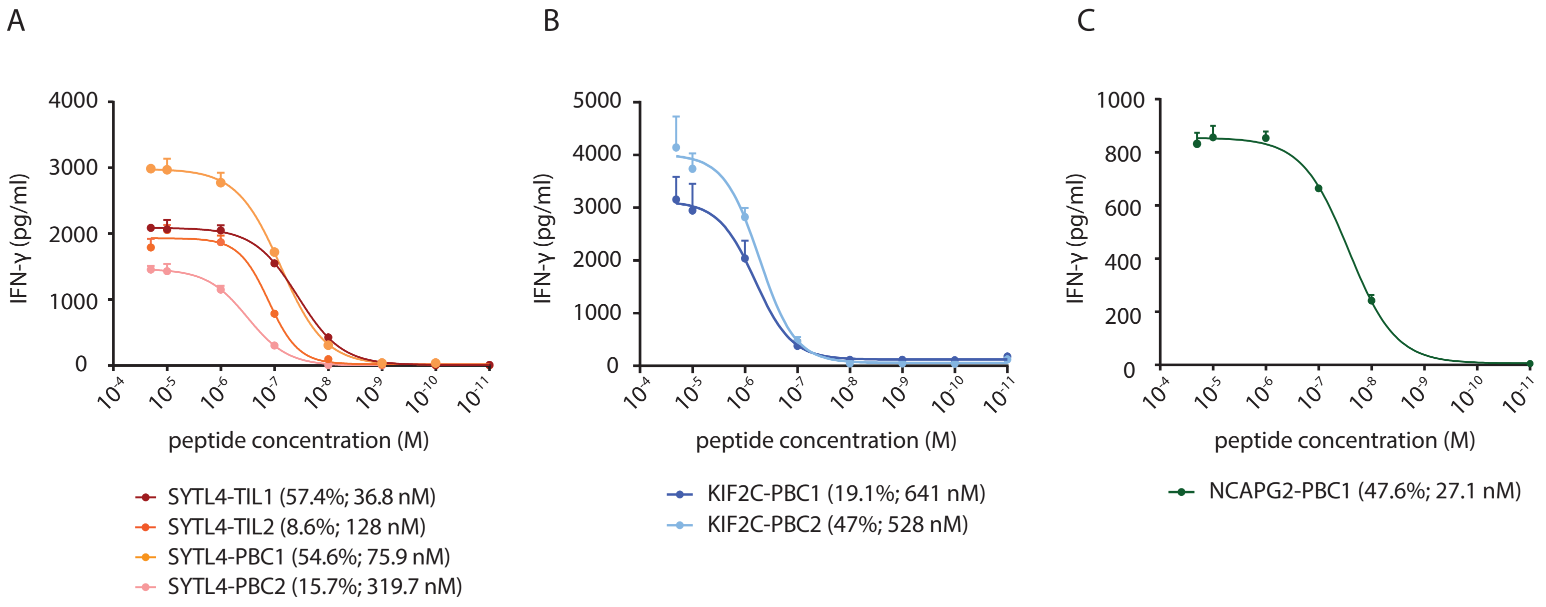
A
B
C
IFN-γ (pg/ml)
peptide concentration (M)
SYTL4-TIL1 (57.4%; 36.8 nM)
SYTL4-TIL2 (8.6%; 128 nM)
SYTL4-PBC1 (54.6%; 75.9 nM)
SYTL4-PBC2 (15.7%; 319.7 nM)
KIF2C-PBC1 (19.1%; 641 nM)
KIF2C-PBC2 (47%; 528 nM)
NCAPG2-PBC1 (47.6%; 27.1 nM)

Figure S6

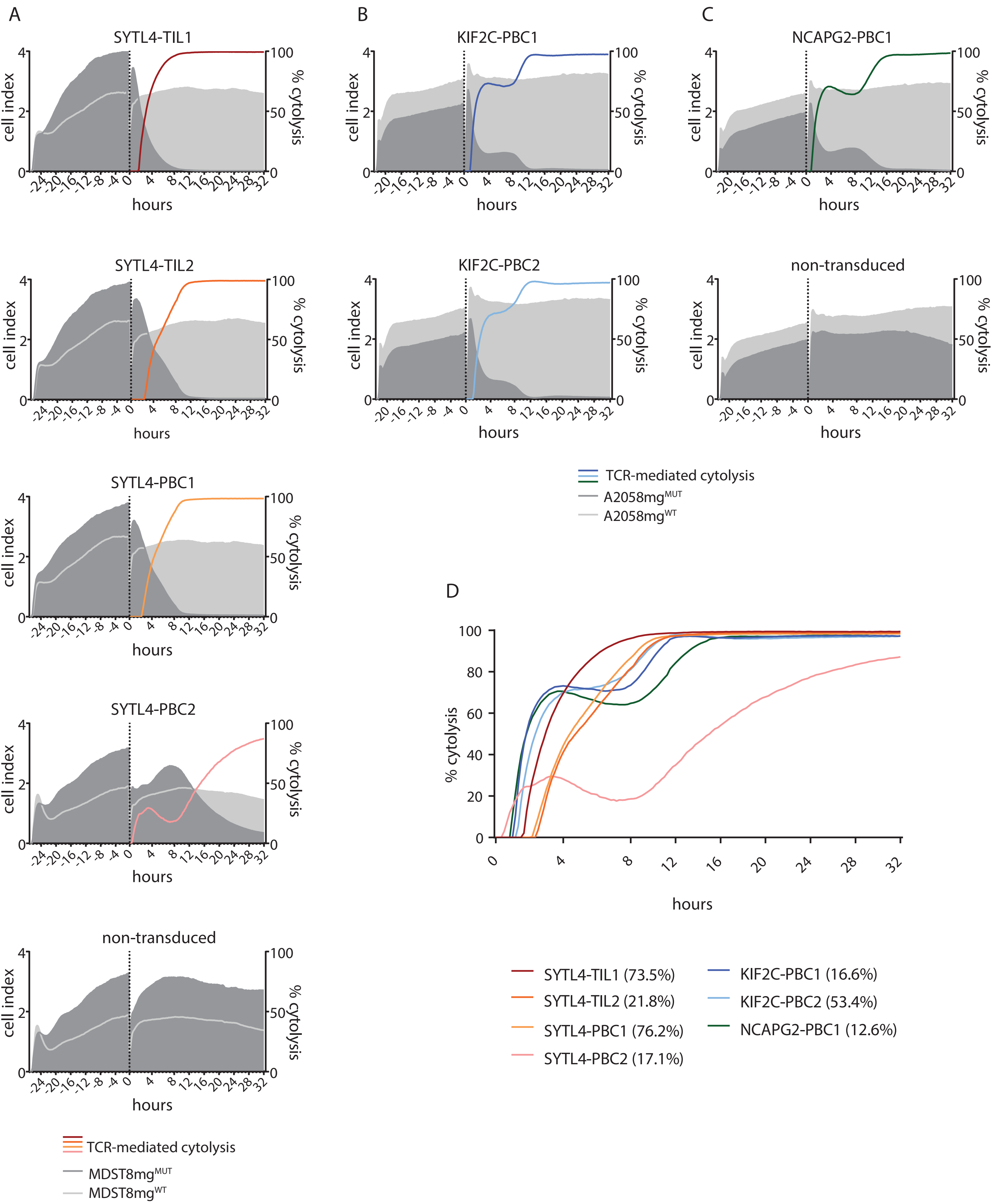
A
SYTL4-TIL1
SYTL4-TIL2
SYTL4-PBC1
SYTL4-PBC2
non-transduced
cell index
% cytolysis
hours
TCR-mediated cytolysis
MDST8mg^MUT
MDST8mg^WT
B
KIF2C-PBC1
KIF2C-PBC2
C
NCAPG2-PBC1
non-transduced
TCR-mediated cytolysis
A2058mg^MUT
A2058mg^WT
D
% cytolysis
hours
SYTL4-TIL1 (73.5%)
SYTL4-TIL2 (21.8%)
SYTL4-PBC1 (76.2%)
SYTL4-PBC2 (17.1%)
KIF2C-PBC1 (16.6%)
KIF2C-PBC2 (53.4%)
NCAPG2-PBC1 (12.6%)

Figure S7

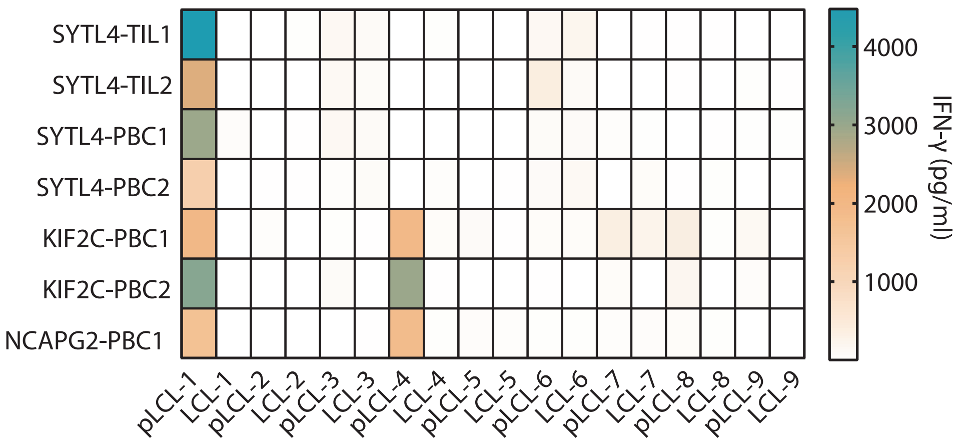
SYTL4-TIL1
SYTL4-TIL2
SYTL4-PBC1
SYTL4-PBC2
KIF2C-PBC1
KIF2C-PBC2
NCAPG2-PBC1
pLCL-1
LCL-1
pLCL-2
LCL-2
pLCL-3
LCL-3
pLCL-4
LCL-4
pLCL-5
LCL-5
pLCL-6
LCL-6
pLCL-7
LCL-7
pLCL-8
LCL-8
pLCL-9
LCL-9
4000
3000
2000
1000
IFN-γ (pg/ml)

Figure S8

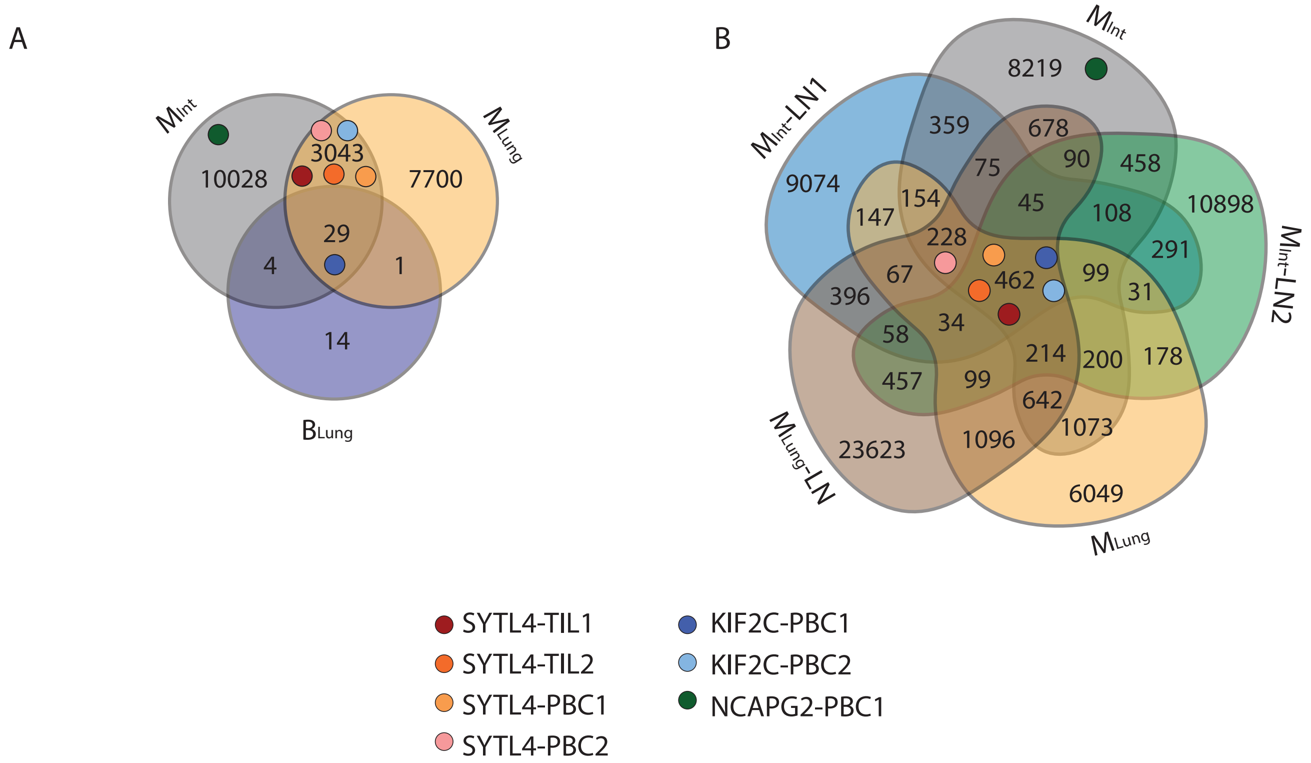
A
B
MInt
MLung
BLung
10028
3043
7700
29
4
1
14
MInt-LN1
MInt-LN2
MLung-LN
8219
359
678
75
90
458
9074
147
154
45
108
10898
228
291
67
462
99
31
396
34
58
214
200
178
457
99
642
1073
23623
1096
6049
SYTL4-TIL1
SYTL4-TIL2
SYTL4-PBC1
SYTL4-PBC2
KIF2C-PBC1
KIF2C-PBC2
NCAPG2-PBC1

Figure S9

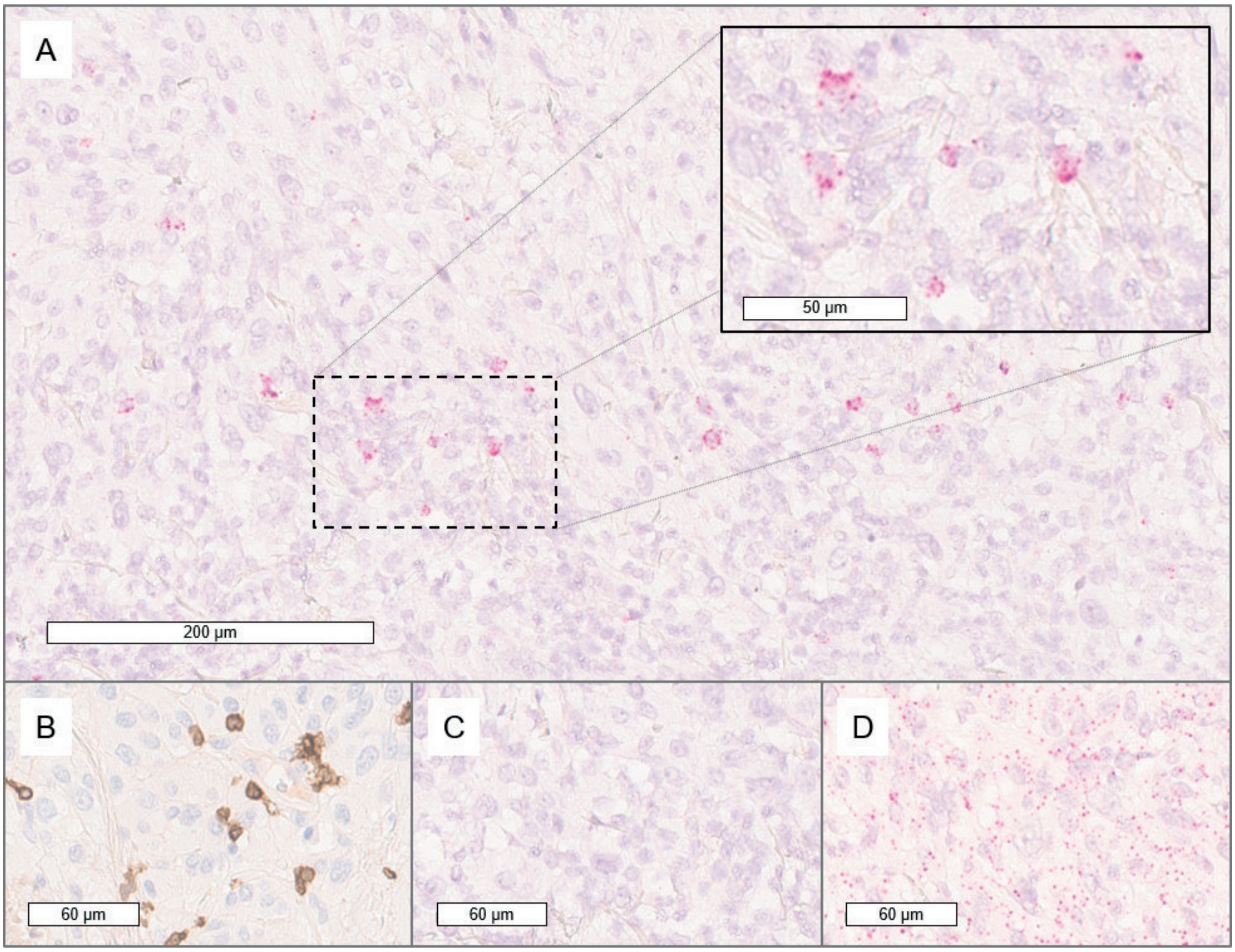


Figure S10

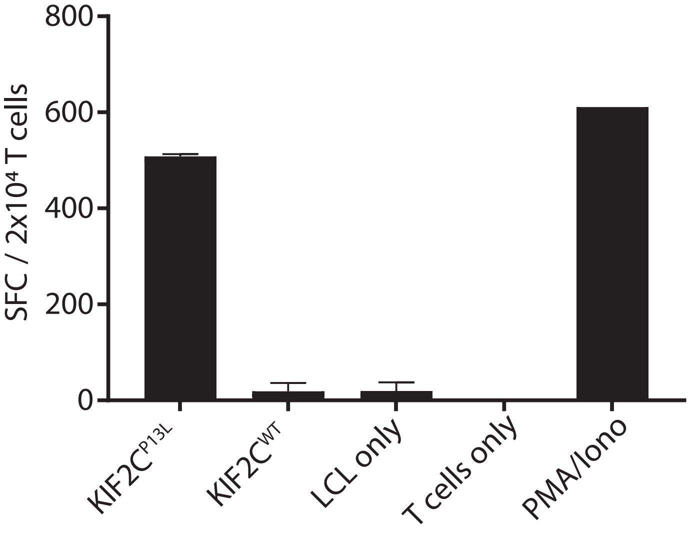

800
600
400
200
0
SFC / 2x10⁴ T cells
KIF2C^P13L
KIF2C^WT
LCL only
T cells only
PMA/Iono


Figure S11

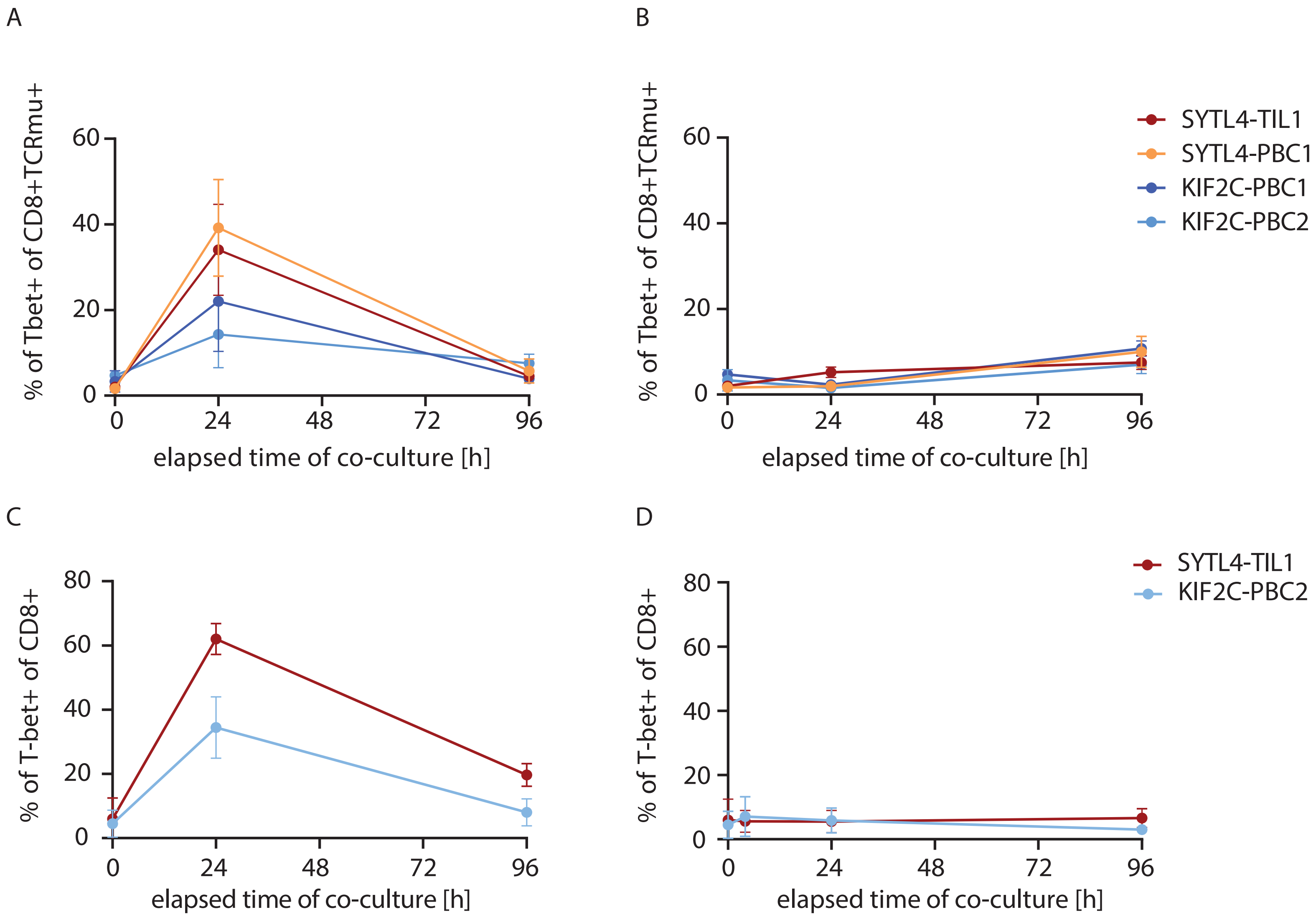
A
B
C
D
% of Tbet+ of CD8+TCRmu+
% of T-bet+ of CD8+
elapsed time of co-culture [h]
SYTL4-TIL1
SYTL4-PBC1
KIF2C-PBC1
KIF2C-PBC2
SYTL4-TIL1
KIF2C-PBC2

Figure S12
